# Ribosome rescue factor PELOTA modulates translation start site choice and protein isoform levels of transcription factor C/EBPα

**DOI:** 10.1101/2023.01.16.524343

**Authors:** Samantha G. Fernandez, Lucas Ferguson, Nicholas T. Ingolia

## Abstract

Translation initiation at alternative start sites can dynamically control the synthesis of two or more functionally distinct protein isoforms from a single mRNA. Alternate isoforms of the hematopoietic transcription factor CCAAT-enhancer binding protein *α* (C/EBP*α*) produced from different start sites exert opposing effects during myeloid cell development. This alternative initiation depends on sequence features of the *CEBPA* transcript, including a regulatory upstream open reading frame (uORF), but the molecular basis is not fully understood. Here we identify *trans*-acting factors that affect C/EBP*α* isoform choice using a sensitive and quantitative two-color fluorescence reporter coupled with CRISPRi screening. Our screen uncovered a role for the ribosome rescue factor PELOTA (PELO) in promoting expression of the longer C/EBP*α* isoform, by directly removing inhibitory unrecycled ribosomes and through indirect effects mediated by the mechanistic target of rapamycin (mTOR) kinase. Our work provides further mechanistic insights into coupling between ribosome recycling and translation reinitiation in regulation of a key transcription factor, with implications for normal hematopoiesis and leukemiagenesis.

## Introduction

The regulation of translation initiation shapes cellular proteomes in organisms ranging from bacteria to humans. In eukaryotes, translation preinitiation complexes typically scan unidirectionally from the 5’ end of mRNAs and initiate translation at the first AUG codon ^1,2^. However, *cis*-regulatory sequences present in the 5’ UTR can alter translation start site choice ^3^. Nature has leveraged this flexibility in start site selection to encode multiple alternative protein isoforms on a single transcript ^4–12^. Initiation at alternative start sites can produce isoforms that either gain or lose domains, changing or even inverting protein function. Even more modest N-terminal extensions and truncations can affect the localization and stability of the resulting proteins. Functionally distinct isoforms produced from alternative initiation have wide-ranging consequences from cellular differentiation ^13,14^ and development ^15,16^ to cell cycle regulation ^17^ and innate immune signaling ^18^. The molecular processes that control the choice between translation start sites and thus alternative protein isoforms are not understood, however.

Alternative start sites also initiate translation of short, upstream open reading frames (uORFs)— ubiquitous regulatory elements present in roughly half of all mammalian transcript leaders ^19,20^. The translation of uORFs interferes with ribosomes reaching downstream coding sequences (CDSes), thereby repressing their translation ^19,21–23^. Productive translation of uORF-containing mRNAs requires either that ribosomes bypass the uORF start codon, in a process called leaky scanning, or that unrecycled ribosomal complexes reinitiate after uORF translation to express the downstream ORF ^21,22^. Additionally, one ribosome translating a uORF can block a second initiation complex from scanning past the uORF, and this inhibitory effect is stronger when ribosomes stall on the uORF ^21,22,24–29^. The interplay between scanning, uORF translation, and reinitiation can provide complex, 5’ UTR-encoded regulation. In mammals, translation reinitiation has been best characterized for the stress-responsive bZIP transcription factor cyclic AMP-dependent transcription factor ATF-4 (ATF4), where reintiation after uORF translation control stress-inducible ATF4 synthesis ^30,31^.

Translation of the critical bZIP developmental transcription factors encoded by *CEBPA* and *CEBPB* is likewise regulated by uORFs ^13,14^. In contrast to the three uORFs of *ATF4*, the single uORF of *CEBPA* regulates the synthesis of two distinct, alternative translation isoforms from the same single-exon mRNA ^13,14^. The longer isoform of C/EBP*α* includes the full N-terminal trans-activation domain. In contrast, a shorter isoform of C/EBP*α*, which is initiated from an internal, canonical AUG start codon, produces a truncated isoform that retains the bZIP domain but lacks most of the transactivation domain (Figure 1A) and can act in a dominant-negative manner by blocking long C/EBP*α* binding and transactivation ^32^. The ratio between these isoforms changes during normal development ^33,34^, and they play distinct roles during myeloid lineage commitment ^35^ and regulate different transcriptional targets ^36^. Mutations that reduce long isoform expression are frequently seen in acute myeloid leukemia (AML), including mutations in *CEBPA* itself that occur in 10-15% of these cancers ^32,37^. Even heterozygous mutations abolishing the long isoform are oncogenic, emphasizing that the stoichiometry of these two isoforms must be maintained for proper differentiation.

**Figure 1:**
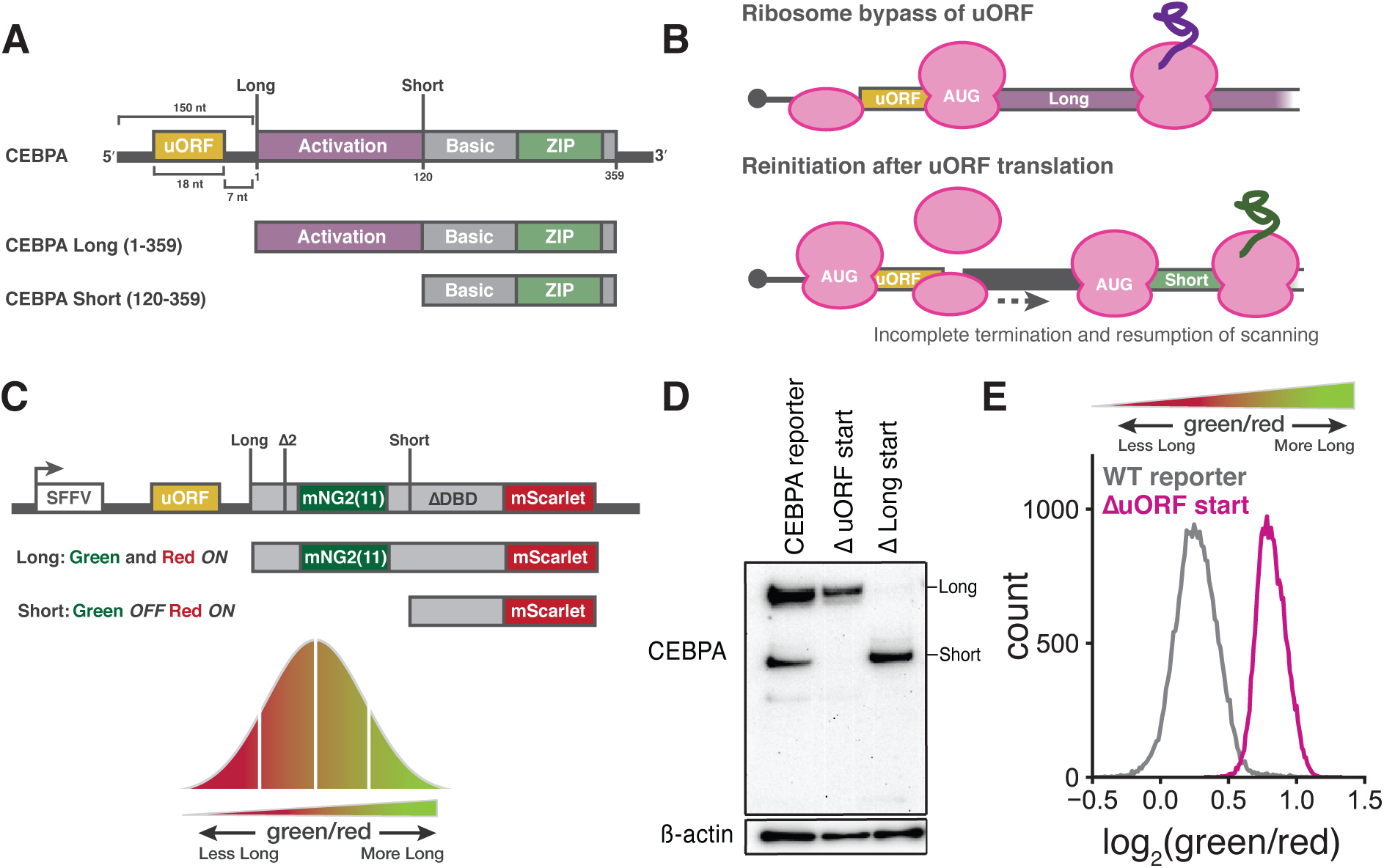
A quantitative, two color fluorescent reporter for measuring start site selection on *CEBPA*. (A) Schematic representation of the *CEBPA* transcript (top) and long and short isoforms (bottom). Length of the 5’ UTR, uORF and intercistronic region are indicated in nucleotides (nt). Amino acid positions of primary start codons and main stop codon are also indicated. (B) Model of long and short isoform translation. Expression of the long isoform relies on leaky scanning past the uORF start codon whereas the short isoform seems to primarily depend on reinitiation after uORF translation. (C) Schematic of the two color reporter assay. Long and short start sites are indicated. The long isoform encodes both green (mNeonGreen2) and red fluorescence (mScarlet) while the short isoform encodes only red fluorescence (mScarlet). Δ2: mutation of a second, in-frame start codon to ACA; Δ*DBD*: deletion of the *CEBPA* basic, DNA-binding domain. (D) Western blot of K562 cells stably expressing CEBPA reporters relative to *β*-actin loading control. Δ uORF start: AUG*→*ACA; Δ Long start: AUG*→*ACA. (E) Flow cytometry measurements of the green/red fluorescent ratio distribution in K562 cells stably expressing either the WT reporter or the Δ uORF start reporter.

This isoform balance is dependent on a short, 6 codon uORF that begins with a canonical AUG start codon and ends just 7 nucleotides before the long isoform start codon ^14^. Translation of the long isoform is mutually exclusive with uORF translation and occurs through leaky scanning past the suboptimal uORF start site followed by initiation at the next start codon. Synthesis of the short isoform, on the other hand, depends on translation of the uORF followed by reinitiation that bypasses the long isoform start codon and instead occurs at a downstream, internal AUG codon (Figure 1B). The molecular details of translation reinitiation are not fully understood, but this process likely involves the ribosome—or at least the small 40S subunit—remaining associated with the mRNA after termination, recruiting a new initiator tRNA, and scanning for another start codon. Few translation initiation factors are known to selectively influence reinitiation; the best characterized reinitiation factors are the density-regulated reinitiation and release factor (DENR) and multiple copies in T-cell lymphoma-1 (MCTS1) complex, first shown to favor reinitiation in *Drosophila melanogaster* ^38^. The DENR/MCTS1 complex selectively facilitates ribosome recycling after uORF translation on the stress-responsive transcription factor *ATF4*, ^39,40^.

Here we characterize the *trans*-regulatory landscape governing translational control of the C/EBP*α* isoform ratio using a dual fluorescent reporter coupled with CRISPRi screening. We identify several factors that influence start site selection on *CEBPA*, including DENR/MCTS1, and uncover a surprising role for the ribosome rescue factor PELO during or after uORF termination. PELO functions in translational quality control pathways to release stalled ribosomes from truncated transcripts and remove unrecycled, post-termination ribosomes from 3’ UTRs or at the ends of truncated transcripts ^41–44^. Our data suggest that PELO enhances long C/EBP*α* isoform expression directly, in addition indirect effects mediated by mTOR activation. Loss of *PELO* may allow post-termination ribosomes to accumulate after uORF translation, and unrecycled ribosomes may provide another layer of uORF-mediated repression.

## Results

### A fluorescent reporter measures *CEBPA* translation start site choice

To measure translation start site selection on *CEBPA*, we developed a dual color reporter that converted the two translational isoforms of C/EBP*α* into two distinct fluorescent proteins produced from the same mRNA. Because the shorter isoform is an N-terminal truncation of the longer isoform, we could not simply mark each isoform with its own fluorescent protein. Instead, we fused the fast-folding, red fluorescent protein mScarlet-I ^45^ to the shared C-terminus, such that it would be expressed in both isoforms. Red fluorescence would therefore serve as a proxy for total protein abundance. We then inserted a green fluorescent protein into the long-isoform-specific N-terminal extension so that green fluorescence reported specifically on long isoform abundance. Thus, the ratio of green to red fluorescence measures the relative abundance of the two translational isoforms (Figure 1C):

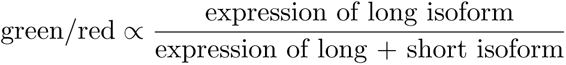

The endogenous N-terminal extension on the long isoform of C/EBP*α* is substantially shorter than a fluorescent protein and its mRNA sequence may contain regulatory information. To minimize disruptions to the organization and regulation of the transcript, we encoded only the short (16 amino acid) fragment of the split, self-complementing green fluorescent protein mNeonGreen2 (mNG2_11_) ^46,47^ in the *CEBPA* CDS between the long and short isoform start sites. Co-expression of our long isoform reporter with the larger fragment of mNeonGreen2 (mNG2_1*−*10_) reconstituted green fluorescence (Figure S1A). To increase the dynamic range and sensitivity of our reporter, we optimized the Kozak sequence around the uORF start codon to enhance short isoform expression. We further deleted the DNA binding domain of C/EBP*α* to mitigate any secondary transcriptional effects of overexpressing our reporter. We then stably integrated a single copy of this construct into the *AAVS1* locus in a K562 human myeloid leukemia cell line stably expressing mNG2_1*−*10_ to generate monoclonal reporter cell lines (Figure S1B).

First, to verify that our reporter recapitulates the regulated choice between *CEBPA* translation start sites, we tested the effects of mutating the start codons in our reporter. Mutation of the uORF start codon abolished short isoform expression by eliminating reinitiation, and mutation of the long isoform start codon itself eliminated long isoform expression (Figure 1D). Furthermore, the reporter with a mutation in the uORF start site—and thus exclusively long isoform expression— showed a much higher ratio of green to red fluorescence relative to the wild-type (WT) reporter, confirming that fluorescence could be used to accurately monitor the isoform ratio (Figure 1E). We further tested how fluorescence of the wild-type reporter changed under conditions that shift the *CEBPA* isoform ratio. Treatment with the allosteric mTOR inhibitor rapamycin reduces short isoform expression ^14^. We recapitulated this effect in our system by treating our reporter cell line with the mTOR active-site inhibitor PP242^48,49^ and observed an increase in the ratio between green and red fluorescence, relative to DMSO-treated cells, consistent with a shift towards long over short isoform expression (Figure S1C). Thus, our fluorescent reporter measures changes in isoform ratio resulting from the choice between translation start sites.

### CRISPRi screens identify genes that modulate *CEBPA* start site choice

We then set out to to identify factors that modulate translation start site selection on our fluorescent *CEBPA* reporter, using CRISPR-based screening. Genetic perturbations that shift the balance between long and short isoforms will change the green/red fluorescence ratio, similar to the effect we saw from mutating the uORF start codon. It is thus possible to select for these perturbations by fluorescence-activated cell sorting (FACS) ^50,51^.

Because many translation factors are essential and mutants can provoke strong growth defects, we perturbed gene expression by CRISPR interference (CRISPRi) ^52^, which produces strong partial loss-of-function phenotypes that allow uniform comparisons across essential and non-essential genes. We transduced our reporter cell line with four different, lentiviral CRISPRi sgRNA sublibraries that collectively comprised 57,900 guides targeting approximately 10,000 genes, in addition to 1,070 nontargeting control guides ^53^. We carried out FACS to distribute transduced cells into four distinct bins depending on their green/red fluorescence ratio and quantified the relative frequency of each sgRNA across the four bins by high throughput sequencing (Figure 2A). Cells expressing a CRISPRi sgRNA that alters the isoform ratio should be unequally distributed across these four FACS bins (Figure S2A).

**Figure 2:**
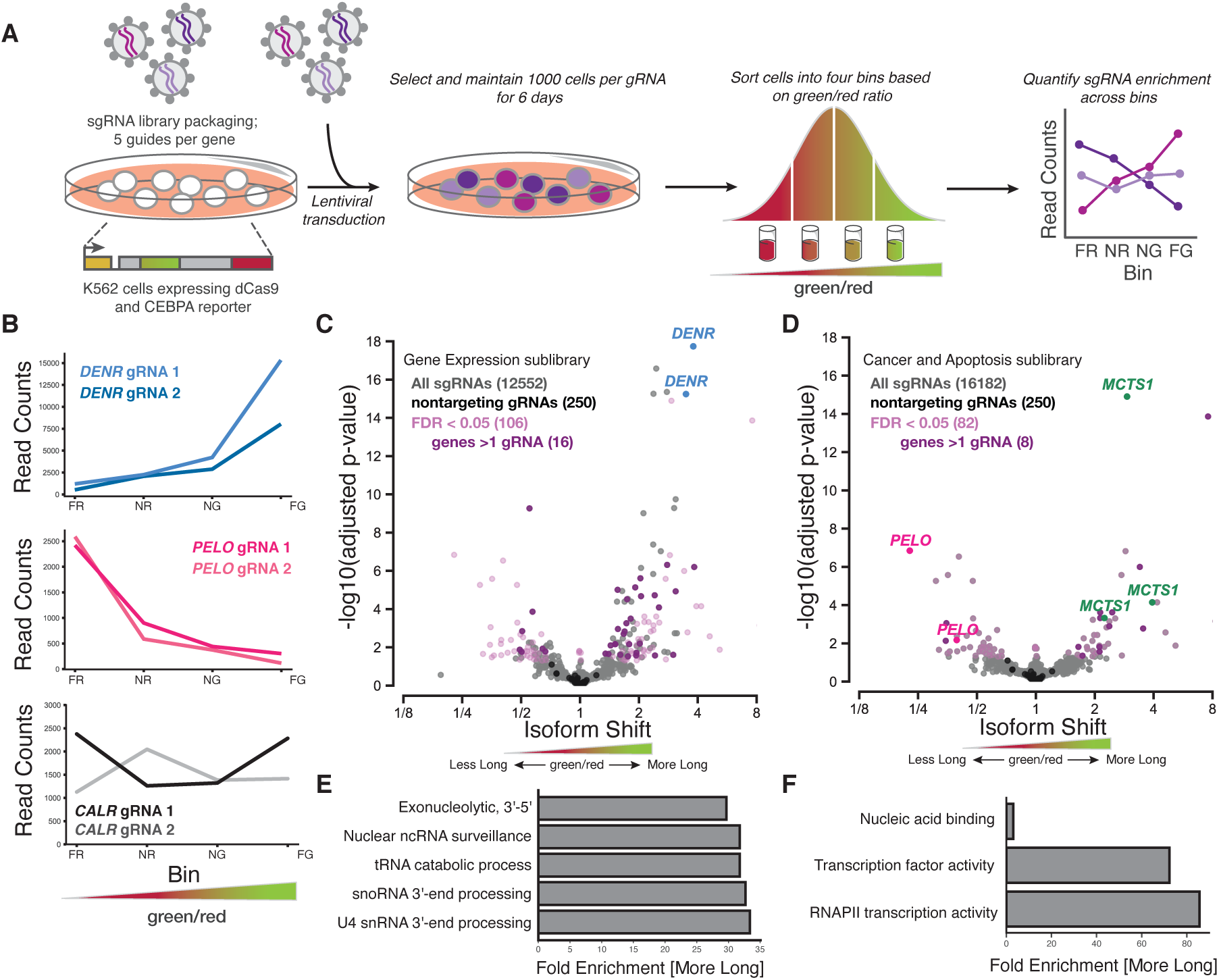
CRISPRi sublibrary screens identify regulators of *CEBPA* isoform expression. (A) Schematic representation of FACS-based CRISPRi screening strategy. Bin labels reflect green/red distribution. FR: far red; NR: near red; NG: near green; FG: far green. (B) Distribution of sgRNA read counts across FACS bins for sgRNAs (in C and D) against *DENR*, *PELO* and *CALR*. (C) Gene Expression sgRNA sublibrary profile representing the relative shift in long and short isoform usage. Each point represents a single sgRNA with sgRNAs against *DENR* highlighted. Colors indicate cutoffs for significance (false discovery rate, FDR *<* 0.05), genes with at least 2 sgRNAs and nontargeting sgRNAs. (D) Cancer and Apoptosis sgRNA sublibrary profile, as in (C). sgRNAs against *MCTS1* and *PELO* are highlighted. (E) Gene ontology (GO) terms associated with sgRNAs that were enriched in cells expressing more long isoform (a higher green/red ratio) in the Gene Expression sublibrary screen. Only sgRNAs with a FDR *<* 0.05 were included in the analysis. GO term analysis was performed using Fisher’s exact test using the Bonferroni correction for multiple testing. Categories chosen represent the most statistically significant terms with a fold enrichment *>* 29. (F) As in (E), but with sgRNAs from the Cancer and Apoptosis sublibrary screen.

Indeed, we identified dozens of sgRNAs that changed the C/EBP*α* isoform ratio. These sgRNAs were strongly shifted towards one side of the sorted population, despite the generally strong correlation in sgRNA abundance across different bins (Spearman’s *ρ* = 0.75 *−* 0.76) (Figures 2B and S2B-D). We quantified the shift in isoform ratio for each sgRNA using a generalized linear model of its abundance in the four sorted bins (Figures 2C and 2D). The vast majority of our 1,070 nontargeting sgRNAs showed no significant shift, demonstrating the specificity of our experimental design and analysis strategy (Figure 2C, 2D, and S2B). Often, two or more independent sgRNAs targeting the same gene caused significant shifts, further arguing that out approach was robust.

Many translation factors emerged among the targets with the strongest and most significant changes in isoform ratio. These included sgRNAs against *DENR* and *MCTS1*, which greatly increased the green/red ratio (Figures 2B-D and S2C) ^54–57^. Loss of *DENR* or *MCTS1* reduces reinitiation after uORF translation in flies ^38^, although work in human cells suggested that they primarily affect transcripts with extremely short, single-codon uORFs ^58^. The shift towards long isoform production in these knockdowns is consistent with a defect in translation reinitiation at the short isoform start codon. Depletion of the *eIF4G* paralog *eIF4G2/DAP5*, also implicated in reinitiation ^59,60^, caused a strong increase in green fluorescence as well (Figure S3A). Interestingly, depletion of the initiation factor eIF2*α*, encoded by *EIF2S1*, actually decreased the green/red ratio, in contrast with the observation that increasing eIF2*α* availability causes a shift towards short isoform production ^14^. More broadly, the targets that increased the long isoform fraction across our screen were enriched for gene ontology (GO) annotations for nucleic acid binding and various mRNA-related regulatory processes (Figure 2E and F).

Depletion of the ribosome rescue factor *PELO* strongly reduced the green/red ratio, indicating a major shift towards short isoform production (Figures 2B, 2D, and S2C). Notably, although *PELO* functions in translation ^43,61,62^, it was not previously linked to uORF-mediated regulation, suggesting a distinct and perhaps *CEBPA*-specific role. Furthermore, *PELO* is homologous to the peptide release factor encoded by *ETF1* that acts in normal translation termination ^63^, and two sgRNAs targeting *ETF1* showed a similar but weaker shift towards short isoform translation (Figure S3A). These results suggested that termination and recycling—perhaps after uORF translation— could affect start site choice and thus C/EBP*α* isoform ratios.

### Reinitiation and ribosome rescue factors control *CEBPA* start site choice

We selected several genes with known roles in translation and RNA biology for individual validation. Two independent, clonal cell lines expressing an sgRNA against *DENR* both had higher green/red fluorescence ratio than cells expressing a nontargeting control sgRNA, in agreement with our results from flow sorting and sequencing; we saw similar results in two independent clones expressing sgRNAs against *DAP5* (Figures 3A and S4A). We also recapitulated a lower green/red ratio in two clonal cell lines expressing an sgRNA against *PELO* (Figures 3A and S4A). More broadly, we saw a strong correlation (*r* = 0.96) between flow cytometry measurements and FACS enrichment across a collection of seven other sgRNAs, each of which shifted the green/red ratio in the direction expected from screening results (Figure 3B).

**Figure 3:**
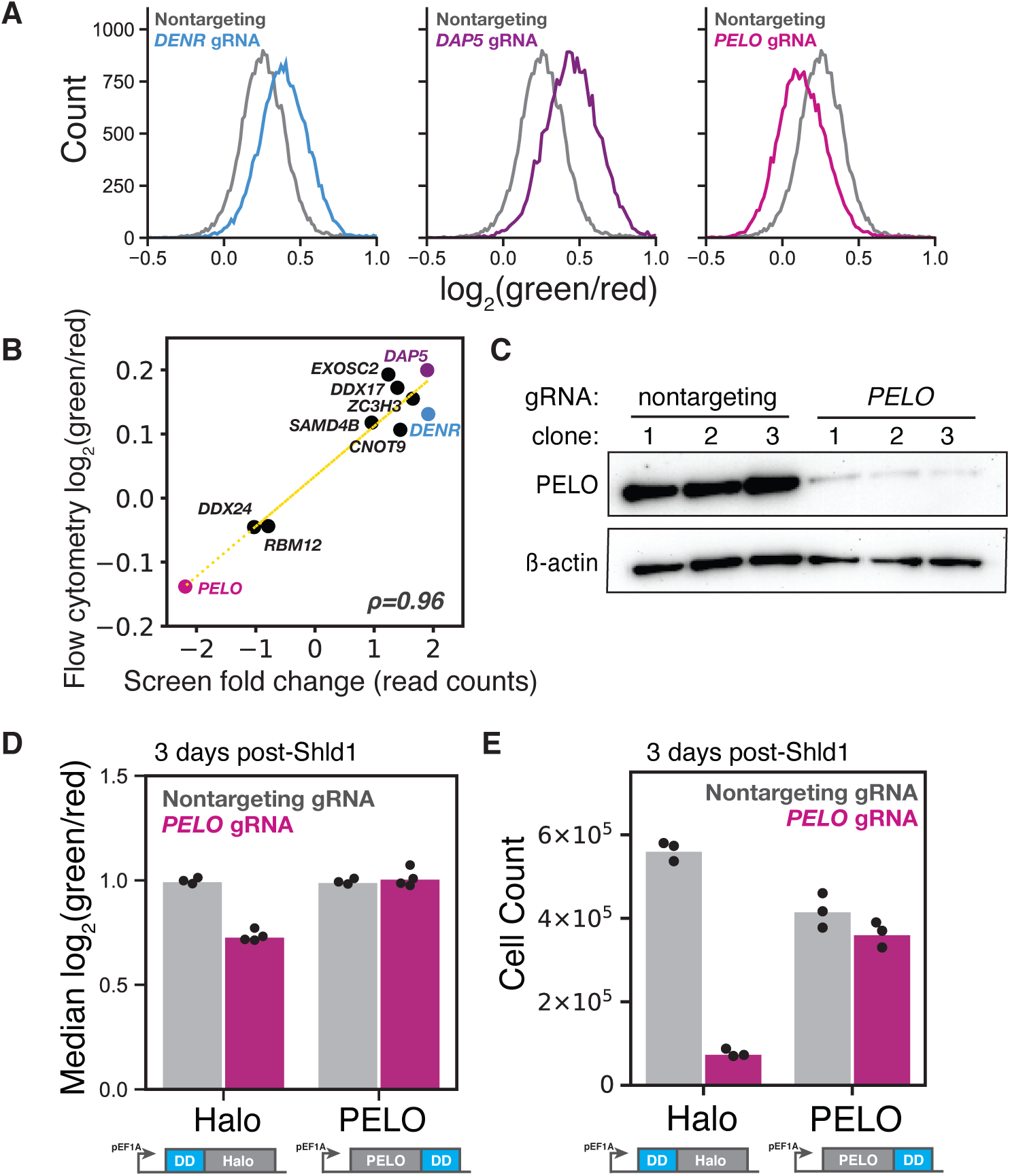
Validation of top CRISPRi sublibrary screen targets. (A) Flow cytometry measurements of the *CEBPA* two color reporter cell line transduced with either a nontargeting sgRNA or the top scoring sgRNAs against *DENR*, *DAP5* and *PELO*. (B) Comparison of the difference in green/red fluorescence ratio by flow cytometry between each target sgRNA and a nontargeting sgRNA and their corresponding read count enrichment in the CRISPRi sublibrary screens, *ρ*=0.96. (C) Western blot of PELO levels relative to *β*-actin in two color reporter lines transduced with either a nontargeting sgRNA or the top scoring *PELO* sgRNA. Clones represent separate sgRNA transductions. (D) Median green/red fluorescence measurements of the two color reporter by flow cytometry in the PELO rescue assay, n = 3 or 4. Re-expression of constructs was induced by 1 *µ*M Shield1 (Shld1) and harvested 72h later. Measurements were normalized to uninduced, nontargeting sgRNA conditions. (E) Median cell count measurements of cell lines in the PELO rescue assay 72h post-Shld1 treament, n = 3.

Short isoform expression depends primarily on downstream reinitiation after uORF translation. In general, ribosomes must be recycled after translation termination in order to prepare them for a new round of initiation. Previous work has shown that the DENR/MCTS1 heterodimer can promote recycling of 40S ribosomes *in vitro* ^64^. More recently, this recycling activity has been linked with reinitiation after uORF translation; DENR/MCTS1 appear to remove tRNAs from post-termination ribosomal complexes, thereby allowing 40S ribosomes to recruit new initiator tRNA and continue scanning ^39,65^. This role in post-termination recycling explains the effects of *DENR* depletion on *CEBPA* start site selection.

The close connection between recycling and reinitiation also suggests how PELO—which is implicated in ribosome rescue and recycling after aberrant termination—could affect the choice between start sites. We then set out to further investigate the link between rescue, recycling and reinitiation. We first confirmed that the isoform shift we saw with sgRNAs targeting *PELO* indeed arose due to PELO depletion. The strongest sgRNA against *PELO* greatly reduced PELO protein levels (Figure 3C). To rescue this PELO depletion, we overexpressed *PELO* from the strong, constitutive *EEF1A* promoter. The rescue construct also contained a drug-controlled destabilization domain (DD) that is stabilized in the presence of the small molecule Shield1 (Figure S5A) ^66^. We stably integrated either the *PELO* rescue construct, or an inactive HaloTag control, into our reporter cell line (Figure S5B). Next, we transduced either a strong *PELO* sgRNA or a nontargeting control into each of these two cell lines and treated cells with Shield1. We found that *PELO* overexpression—but not HaloTag overexpression—completely rescued the *PELO* knockdown phenotype (Figure 3D) and mitigated the substantial cell viability defect caused by *PELO* depletion as well (Figure 3E). To confirm that the effect of *PELO* was not an artifact of our modified uORF start context, we recapitulated *PELO* knockdown and rescue with a reporter harboring the native Kozak sequence around the uORF start codon (Figures S5C and S5D).

While it seemed likely that *PELO* depletion affected start site selection, we wanted to exclude the possibility that it had isoform-specific, post-translational effects on protein stability. To do so, we engineered a cell line containing a variant *CEBPA* reporter fused to a C-terminal, destabilizing PEST sequence, ensuring the rapid turnover of both isoforms. In this cell line, we again recapitulated the reduction in the green/red fluorescence ratio upon sgRNA-mediated *PELO* knockdown, arguing that this shift was not mediated by differences in protein half-life between the two isoforms (Figure S5E). Instead, it appears that PELO plays an uncharacterized role in regulating translation start site selection on *CEBPA*.

### PELO effects on *CEBPA* translation depend on uORF length

As PELO has not previously been implicated in uORF-mediated regulation, we next asked whether its effect on the relative abundance of the two C/EBP*α* isoforms depended on the uORF. We generated cell lines expressing either a reporter variant with a mutation inactivating the uORF start codon (Figure 1D), or a reporter with a short, unstructured 5’UTR containing no uORFs in place of the endogenous 5’UTR of *CEBPA*. Eliminating the uORF start codon or replacing the whole 5’UTR reduced short isoform expression, as reflected in the higher green/red fluorescence ratio observed in these cell lines (Figure 4A). The effects of *DENR* and *DAP5* knockdown were weakened or eliminated in these two reporters, which should no longer support reinitiation. The effects of *PELO* knockdown were likewise greatly attenuated in variants without uORFs; *PELO* depletion did not reduce the high green/red ratio (Figure 4B). These observations suggest that the presence—and translation—of the *CEBPA* uORF is required for the change in isoform usage induced by PELO on our reporter.

**Figure 4:**
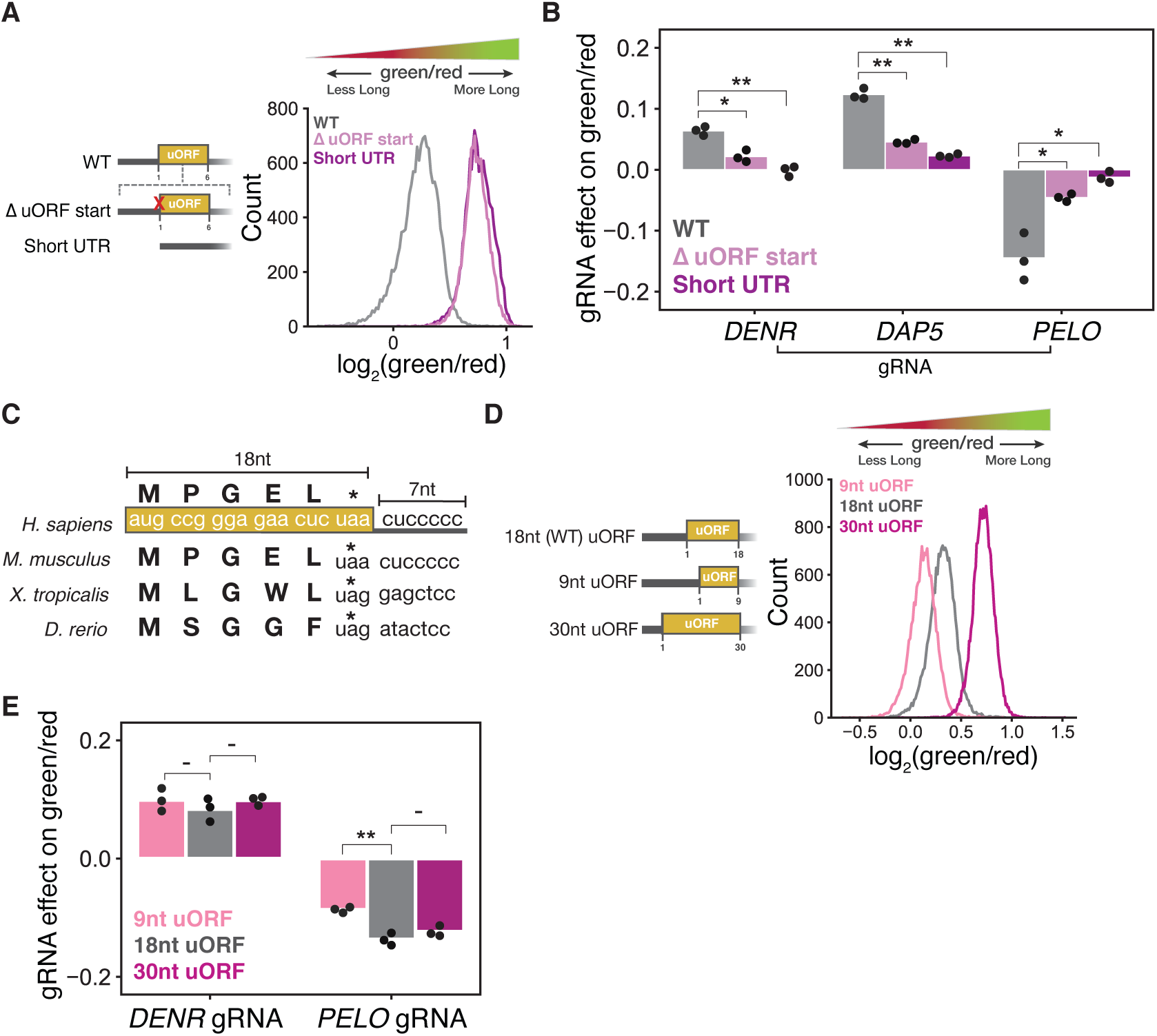
The effect of PELO on *CEBPA* translation depends on the uORF length. (A) Distribution of green/red ratio by flow cytometry of cell lines stably expressing two color reporters with either a Δ uORF start codon mutant or a short (22 nucleotides), unstructured 5’UTR with no uORFs. (B) Difference in median green/red fluorescence ratio by flow cytometry between each target sgRNA and a nontargeting sgRNA in the indicated stable cell lines. Statistical significance was calculated by two-sided t-tests with p-values denoted by: *∗* : *<* 0.05 and *∗∗* : *<* 0.01. (C) Schematic of *CEBPA* uORF with length of uORF and intercistronic region indicated in nucleotides (nt). (D) Distribution of green/red ratio by flow cytometry of cell lines stably expressing two color reporters with varying uORF lengths. Length of uORFs are indicated in nucleotides (nt). The distance between the uORF stop codon and main start codon is 7 nt in all reporters. (E) Median difference in green/red fluorescence ratio by flow cytometry between each target sgRNA and a nontargeting sgRNA in the indicated stable cell lines. Statistical significance was calculated as in (B).

We next considered the distinctive features of the *CEBPA* uORF. The length of the uORF and the short, 7 nucleotide separation between the end of the uORF and the long isoform start codon are conserved across *CEBPA* homologs, although their sequence varies (Figure 4C). We thus tested whether uORF length impacted the effect of our knockdowns by generating stable two color reporter cell lines encoding the *CEBPA* uORF with varying lengths that still preserved the distance between the uORF stop codon and main CDS start codon. Shortening the uORF to 9 nucleotides reduced the green/red ratio, suggesting a shift towards short isoform production, in line with the general observation that shorter uORFs confer higher reinitiation probability ^67^. In contrast, increasing the length of the uORF to 30 nucleotides increased the green/red ratio, indicating less efficient reinitiation (Figure 4D).

When we introduced sgRNAs targeting *PELO* into each of these reporter lines, we found that the effect of *PELO* knockdown was most diminished on the shortest, 9 nucleotide uORF and was unchanged in the 30 nucleotide uORF reporter relative to the wild type, 18 nucleotide uORF reporter. In contrast, *DENR* knockdown produced a similar effect on the green/red ratio across all reporter lines (Figure 4E). These results suggest that unlike DENR, *PELO* loss is especially sensitive to the relative positioning of the uORF start and stop codons.

### *PELO* knockdown decreases long isoform expression

Our two-color reporter provides a sensitive measure of changes in the isoform ratio, but does not distinguish whether *PELO* knockdown reduces long isoform expression or increases short isoform expression. To delineate between these possibilities, we expressed a third fluorescent protein that would serve as a normalizing control and allow us to quantify the absolute abundance of each *CEBPA* isoform. We chose the infrared fluorescent protein iRFP670^68,69^, which is spectrally distinct from mNeonGreen2 and mScarlet-I, enabling simultaneous quantification of all three fluorescent proteins by flow cytometry, and expressed it using the constitutive *EEF1A* promoter. We calibrated our fluorescence measurements using a reporter with a mutation in the uORF start codon that expresses exclusively the long isoform, as well as a reporter that expresses only the short isoform (Figures 1D and 5A).

**Figure 5:**
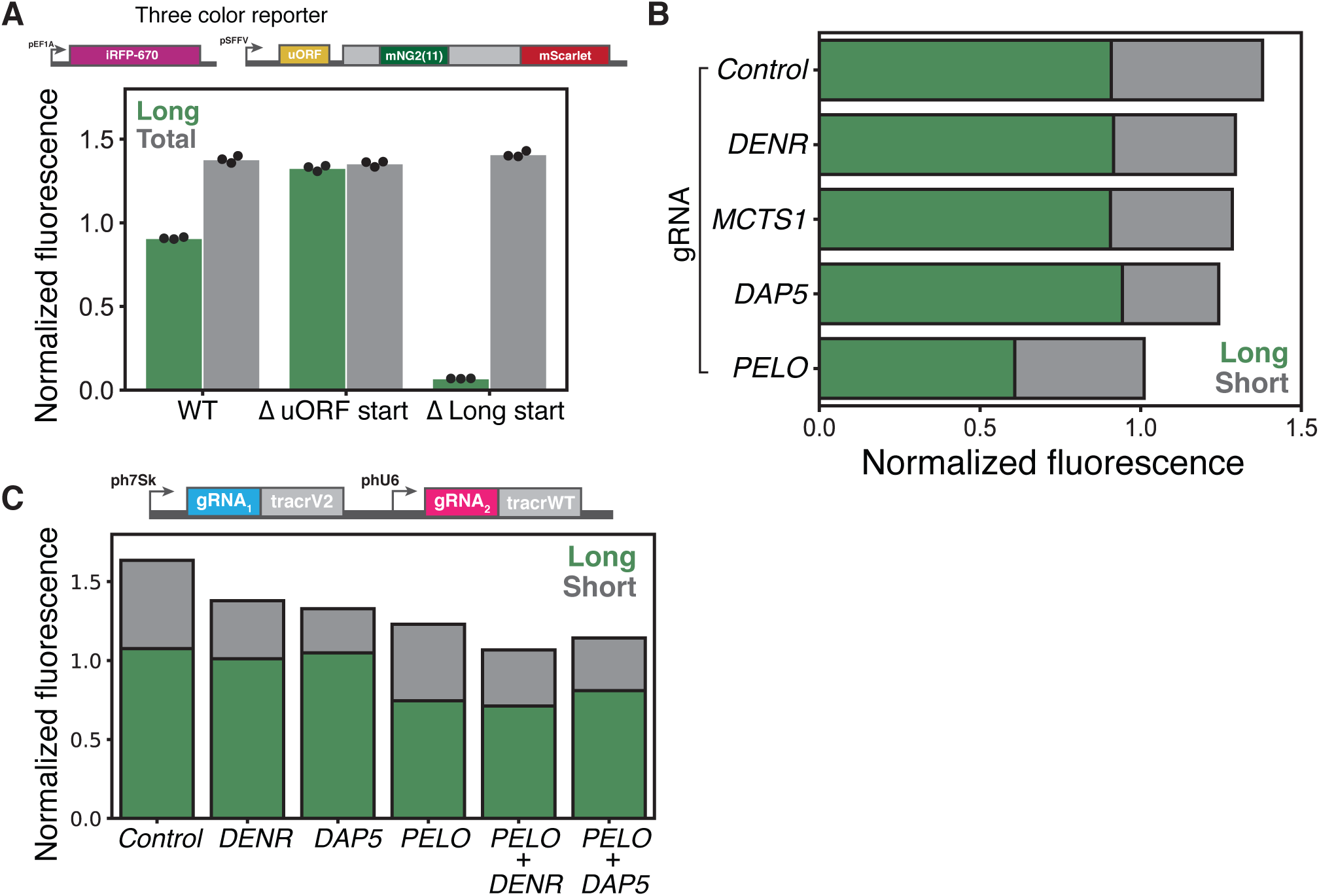
Three color reporter reveals that *PELO* knockdown decreases long isoform expression. (A) Schematic of three color reporter assay (top). Flow cytometry measurements of median green and red fluorescence normalized to IRFP670 in stable cell lines expressing the three color reporter, the Δ uORF start codon mutant reporter or the Δ Long start codon mutant reporter, n = 3 (bottom). (B) Flow cytometry measurements of median green and red fluorescence normalized to IRFP670 in stable cell lines expressing the wild type three color reporter transduced with indicated sgRNAs. (C) Median green and red fluorescence normalized to IRFP670 in stable cell lines expressing the wild type three color reporter transduced with indicated dual sgRNAs.

We then used our calibrated fluorescent measurements to characterize the effect of several sgRNA knockdowns on absolute isoform abundance. We first confirmed that our targeted sgRNA knockdowns did not significantly impact the levels of our iRFP670 normalizer reporter (Figure S6A). Consistent with their roles in promoting reinitiation at the short isoform start codon, depletion of *DENR* or *MCTS1* notably reduced red fluorescence with no significant change in normalized green fluorescence, and thus no difference in long isoform levels. We observed a similar reduction in short isoform expression in *DAP5* knockdown cells, in agreement with its proposed role in reinitiation. In *PELO* knockdown cells, we observed a decrease in long isoform expression while short isoform abundance was largely unaffected (Figures 5B and S6B), implying that PELO normally promotes long isoform expression.

We next investigated the interactions between these distinct effects on long and short isoform translation. We knocked down either *DENR* or *DAP5* in combination with *PELO* and compared these effects with single gene knockdown using a dual-sgRNA expression vector ^70^ (Table S1). The effect of depleting both *DENR* and *PELO* was additive—double knockdown reduced short isoform expression to the same extent as *DENR* knockdown alone, and long isoform expression to the same degree as *PELO* single knockdown. Interestingly, depleting *DAP5* and *PELO* together somewhat weakened both the *PELO*-dependent loss of the long isoform as well as the *DAP5*-dependent loss of short isoform expression (Figures 5C and S6C). Nonetheless, the lack of strong epistasis argues that PELO is not directly affecting reinitiation.

### *PELO* depletion increases *CEBPA* uORF translation and activates mTOR

To directly measure the translational effects of *PELO* depletion on our reporter, and across the transcriptome, we performed ribosome profiling ^71,72^ in our reporter cell line transduced with either the top scoring *PELO* sgRNA or a nontargeting sgRNA. Biological triplicates of the same sgRNA treatment correlated very well (*p >* 0.99) and showed clear, sgRNA-specific differences (Figures S7C and S7D). We saw the characteristic accumulation of footprints in the 3’ UTR in our *PELO* knockdown that correspond to unrecycled, vacant 80S ribosomes (Figure S7A), as has been previously reported in both yeast and humans ^43,44^. We further observed a striking increase in ribosome occupancy in the 5’ UTR of our *CEBPA* reporter in *PELO* depleted cells relative to our control. These included footprints on the *CEBPA* uORF, indicative of increased uORF translation in *PELO* knockdown. We also observed a surprising accumulation of footprints that mapped to the long isoform start codon. While these footprints could be derived from ribosomes initiating at the long isoform start codon, our fluorescence measurements indicate that *PELO* depletion reduces long isoform production. Alternately, these footprints could originate from vacant ribosomes that accumulate after uORF termination, analogous to the unrecycled ribosomes that are enriched in 3’ UTRs after main ORF termination as a consequence of reduced PELO levels. Greater persistence of these unrecycled ribosomes could occlude the long isoform start codon in *PELO* knockdown. They could also enhance uORF translation by stalling scanning, pre-initiation complexes at the uORF start codon, leading to a self-reinforcing situation where ribosomes that terminate after uORF translation favor subsequent uORF translation rather than long isoform translation. As this self-reinforcing effect depends on the precise position of the post-termination ribosome relative to the uORF start codon, it also explains the uORF length dependency of the *PELO* knockdown phenotype (Figure 4E). Overall, these results support a model in which *PELO* loss leads to enhanced uORF translation at the expense of long isoform expression (Figures 6A and S7B).

**Figure 6:**
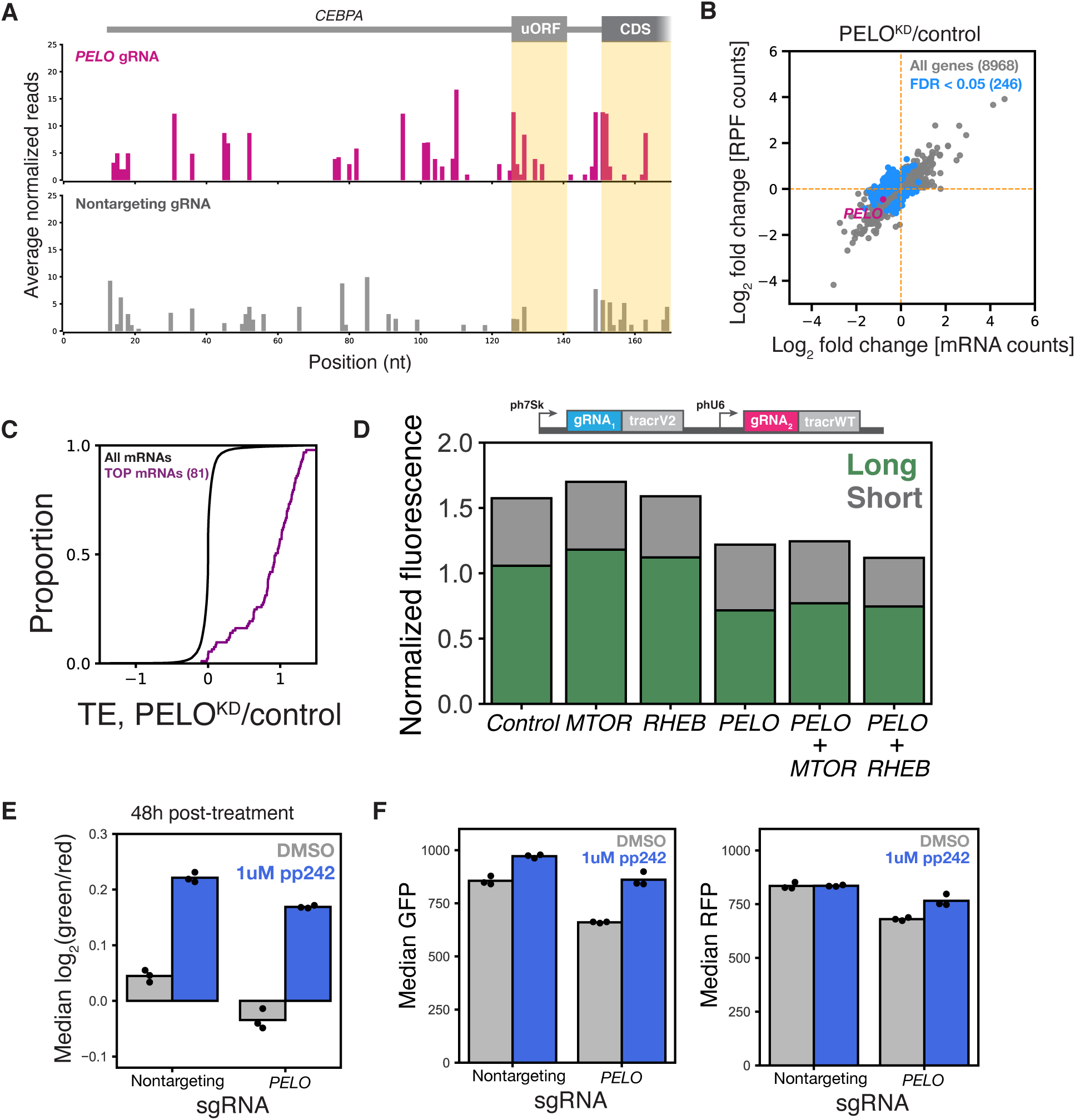
*PELO* knockdown increases uORF translation and activates mTOR. (A) Ribosome occupancy profile of *CEBPA* 5’ UTR. Start and stop codons belonging to the uORF and long isoform start codon are indicated. Read counts were normalized by CDS occupancy and the median read count was determined across replicates, n=3. (B) Scatterplot of log_2_ fold changes (*PELO* KD/control) in ribosome footprint and RNA-seq counts. Each point represents a single gene. Genes with a statistically significant (*F DR <* 0.05) log_2_ fold change in translation efficiency (TE) are indicated in blue. (C) Cumulative distribution of TE log_2_ fold changes of TOP-containing mRNAs. (D) Median green and red fluorescence normalized to IRFP670 in stable cell lines expressing the wild type three color reporter transduced with indicated dual sgRNAs. (E) Median green/red fluorescence measurements in reporter cell lines transduced with the indicated sgRNAs. Cells were treated with either DMSO or 1 *µ*M PP242 (Sigma-Aldrich) 6 days post-transduction. Cells were harvested and assayed by flow cytometry 48h post-drug treatment. (F) Flow cytometry measurements of median green and red fluorescence from (E), n = 3.

In addition to these *CEBPA*-specific effects, we observed a number of translational changes across the transcriptome. We computed translation efficiency (TE) as the ratio of ribosome footprint abundance to matched mRNA abundance. Overall, 248 genes displayed a significant (FDR-adjusted *p* ≤ 0.05) TE difference in our *PELO* depletion (Figure 6B). Among the genes with the strongest increase in TE, 32% (81 genes) were 5’ terminal oligopyrimidine (TOP) motif containing mRNAs ^73^ primarily encoding ribosomal and ribosome-associated proteins (Figure 6C). Translational upregulation of 5’TOP mRNAs is a hallmark of mTOR activation. Indeed, previous work in human fibroblasts and in mouse models also observed a marked translational enhancement of mTOR regulated transcripts in *PELO* knock-outs ^74,75^. mTOR activation has broad ranging effects on protein synthesis and acts primarily on translation initiation by altering availability of the cap-binding protein eIF4E ^76,77^. In fact, previous work has shown that rapamycin-induced mTOR inhibition favors C/EBP*α* long isoform expression by decreasing eIF4E availability ^14^, raising the possibility that mTOR activation explains the reduced long isoform expression in *PELO* knockdown. We thus wanted to ask whether *PELO* affects C/EBP*α* translation above and beyond mTOR-mediated changes.

Despite the known effect of mTOR activity on *CEBPA* translation, genes from this pathway did not stand out in our screen. We identified a modest but significant effect from just one sgRNA targeting *MTOR* itself, and no significant effects from sgRNAs targeting mTOR regulators *RHEB* or *TSC1* (Figure S7E). Individual knockdown of *MTOR* led to a slight increase in the green/red ratio, much smaller than the change seen in *PELO* knockdown, consistent with the weak phenotype in our screening data. We likewise saw that targeted knockdown of the mTORC1 activator *RHEB* did not change the fluorescence phenotype (Figures 6D and S7E), nor did an individual sgRNA against the negative mTOR regulator *TSC1* (Figures S7E and S7F).

CRISPRi knockdown produces only partial loss of function phenotypes, and mTOR activity is modulated by feedback at many levels that might buffer the effects of genetic perturbation. We thus wanted to confirm that *MTOR* knockdown reduced phosphorylation of mTOR targets. Indeed, CRISPRi against *MTOR* reduced phosphorylation of 4EBP1 and almost abolished phosphorylation of RPS6, an indirect translation-related target, confirming that *MTOR* knockdown reduces mTOR activity (Figure S7G). We also observed that *PELO* knockdown increases mTOR activity (Figure S7G).

We then depleted *PELO* in conjunction with either *MTOR* or the mTORC1 activator *RHEB*. Critically, the *PELO* depletion phenotype was unaffected by simultaneous knockdown of either *MTOR* or *RHEB*, arguing that it does not depend on increased mTOR activity. We further tested how the *PELO* knockdown phenotype was affected by the potent mTOR active-site inhibitor PP242, which has a stronger effect than *MTOR* knockdown (Figures 6D and S1C). We transduced our reporter line with either a *PELO* sgRNA or a nontargeting control and treated these cell lines with PP242. Consistent with the strong effect of PP242 in other settings, we saw that it substantially increased the green/red ratio in *PELO* knockdown cells (Figure 6E). Nonetheless, *PELO* knockdown still reduced the green/red ratio in the context of PP242 treatment relative to a nontargeting control, consistent with a direct, mTOR-independent role for PELO in start site selection. Further analysis indicated that PP242 treatment alone enhanced long isoform expression at the expense of short isoform synthesis, with little change in total abundance (Figure 6F). When *PELO* was depleted, PP242 treatment enhanced long isoform production but did not restore total protein levels relative to control cells (Figure 6F). This again suggests that lack of *PELO* directly impairs long isoform synthesis—perhaps by physical obstruction by unrecycled ribosomes—an effect that cannot be suppressed by mTOR inhibition.

## Discussion

We survey the *trans*-acting factors that control the choice between alternate translation start sites that produce opposing isoforms of the key hematopoietic transcription factor C/EBP*α*. Known reinitiation factors *DENR*/*MCTS1* and *DAP5* play substantial roles in promoting short isoform expression, supporting uORF-dependent reinitation as the mechanism for translation from the downstream start codon. We also found that loss of ribosome rescue factor *PELO*, which has no described role in translation reinitiation or uORF-mediated regulation, reduces C/EBP*α* long isoform expression. Our work thus reveals an unexpected link between ribosome rescue and uORF-mediated translational regulation. We propose that long isoform initiation is blocked by unrecycled ribosomes that accumulate after uORF translation, providing a mechanistic connection between ribosome rescue and start site selection.

Both PELO and DENR/MCTS1 have established molecular functions at the end of translation. The impact of depleting these factors on *CEBPA* start site choice suggests that the fate of ribosomes after uORF translation controls downstream translation. The DENR/MCTS1 heterodimer recycles 40S subunits after termination, and this activity is important for subsequent reinitiation in many situations, including *ATF4* translation ^38,39,56,58^. The *CEBPA* uORF may rely on *DENR* because it contains a Leu codon in the penultimate codon position, which confers a particularly strong dependency on DENR for post-termination tRNA eviction ^39,64^.

While PELO is also associated with ribosome recycling ^41,42^, loss of *PELO* affects long isoform initiation specifically and so it does not seem to prepare ribosomes for reinitiation. Instead, in keeping with its function in other contexts, PELO likely removes vacant, un-recycled ribosomes that would otherwise impede subsequent rounds of translation initiation on the transcript. The conserved length and position of the *CEBPA* uORF, near the long isoform start codon, is expected to enhance the consequences of a post-termination stall, because an 80S ribosome that is not released after uORF translation would be positioned over the long isoform start codon, thereby blocking its expression. The specific positioning of this stalled post-termination ribosome could also have an additional effect: because the combined 25-nucleotide length of the uORF and intercistronic region is nearly the size of a 28-nucleotide elongating ribosome footprint, incoming pre-initiation scanning complexes would be queued behind the stalled 80S and positioned near the uORF start codon, enhancing uORF translation. We found that a variant reporter with a shorter uORF was less responsive to *PELO* depletion, consistent with a requirement for the conserved spacing of the *CEBPA* uORF to strengthen the effects of unrecycled ribosomes on both uORF and long isoform initiation. This effect of post-termination complexes is reminiscent of other regulatory paradigms where translational stalling promotes upstream initiation ^27,78–80^.

The *CEBPA* 5’ UTR is highly structured in addition to containing a uORF, making it a prime target for *DAP5* dependence as well ^59^. While the molecular details of DAP5 function are not clear, its general role in downstream CDS translation in the context of structured 5’ UTRs is consistent with its importance for short isoform translation on *CEBPA*. It is also notable that *DAP5*-dependent translation requires efficient termination and recycling, again connecting the results of our screen to uORF translation termination.

We also found that *PELO* depletion activates the mTOR pathway, as seen previously in other systems ^74,75^. Changes in mTOR activity have been shown previously to regulate translation start site usage on *CEBPA* ^14^. Through a combination of genetic and chemical perturbations, we provide evidence for both mTOR dependent and independent effects of *PELO* depletion. Indeed, impaired ribosome recycling after *PELO* depletion may reduce translational capacity ^81^, activating mTOR in a compensatory response. The mechanism linking *PELO* depletion with mTOR activation is unknown, but may involve the direct association of mTORC1 with ribosomes ^82^ or an imbalance between ribosome availability and protein biosynthesis capacity ^81^.

Ribosome rescue and recycling activity are themselves dynamically regulated during erythroid development ^44^. In both hematopoietic progenitor CD34+ cells and K562 cells, PELO is initially upregulated then gradually decreases during differentiation; PELO levels are greatly diminished in primary platelets and reticulocytes relative to proliferative, nucleated cells. Platelets and reticulocytes derive from the common myeloid progenitor, where CEBPA plays a central role in fate specification. In addition to erythroid and megakaryocyte/platelet lineages, the common myeloid progenitor gives rise to the granulocyte/monocyte progenitor, and CEBPA is required specifically for this cell fate decision ^83^. The long isoform of CEBPA is critical for both granulocytic and monocytic maturation while overexpression of the short isoform blocks granulocyte differentiation ^84^. Variations in PELO levels in these blood cell lineages may affect *CEBPA* isoform balance and by extension, myeloid cell fate commitment. Expression of *PELO* is also decreased in AML ^85^, and our data suggest that this would favor the oncogenic, short isoform. These trends underscore the broader physiological and pathological impact of PELO and the ribosome rescue pathway on the regulation of hematopoeisis.

## Acknowledgements

We thank J. Wren Kim, Liana Lareau, and members of the Ingolia and Lareau labs for invaluable, scientific discussions. mNeonGreen2 plasmids were kind gifts of Siyu Feng. We also thank Jonathan Weissman’s lab for the K562 CRISPRi cell line. Hector Nolla, Alma Nuguid Valeros and Kartoosh Heydari provided indispensable support at the Flow Cytometry Facility at UC Berkeley. We also thank the Vincent J. Coates Genomics Sequencing Laboratory at UC Berkeley and the UC Berkeley DNA Sequencing Facility. This work was supported by the National Institutes of Health (www.nih.gov), grants DP2 CA195768 and R01 GM130996 (NTI) and shared instrumentation grant S10 OD018174. The funders had no role in study design, data collection and analysis, decision to publish, or preparation of the manuscript.

## Declaration of Interests

N.T.I. declares equity in Tevard Biosciences and Velia Therapeutics.

## Materials and Methods

### Plasmid and two color reporter construction

All plasmids and primers used are listed in Table S1 and S2, respectively. All plasmids (with the exception of sgRNA expression vectors, see section) were generated by Gibson assembly ^86^ from amplicons made with primers indicated in Table S2.

In brief, to construct the CEBPA two color reporter, the mNG2_11_ sequence was subcloned from a pCMV-mNG2_11_-H2B plasmid kindly gifted by Siyu Feng ^47^ and assembled into codon position 45 in the N-terminus of *CEBPA*. All Met codons in mNG2_11_ were removed to avoid generating new in-frame start sites and substituted with either Val or Ile to preserve hydrophobicity. Similarly, an in-frame start codon at position 14 in *CEBPA* was mutated to ACA (Δ2) to ensure that only the long and short start sites were used. Furthermore, the basic, DNA-binding domain was deleted (Δ*DBD*) to suppress cell proliferation arrest caused by ectopic expression of *CEBPA* ^14,87^ and the Kozak sequence around the uORF start codon was optimized (gccgccATGg, as in Calkhoven et al. ^14^) to increase the dynamic range of our reporter readout. *Hs* CEBPA cDNA was amplified from the genome. Amplicons corresponding to the SFFV promoter, CEBPA 5’UTR, CEBPA-mNG2_11_ and CEBPA-mScarlet-1XFlag were then introduced into the Sbf1 and Kpn1 cut sites of the pNTI620 vector.

### Cell culture

Human K562 cells were grown in RPMI-1640 with L-glutamine (ThermoFisher Scientific) supplemented with 10% FBS, 1% sodium pyruvate, 100 units/mL penicillin and 100 mg/mL streptomycin. Human HEK 293T Lenti-X cells were grown in DMEM + GlutaMax (ThermoFisher Scientific) supplemented with 10% FBS, 1% HEPES, 100 units/mL penicillin and 100 mg/mL streptomycin. K562 cells were maintained at a cell density of 0.5*x*10^6^/mL. All cell lines were obtained from the UC Berkeley Cell Culture Facility and grown at 37*^◦^*C and 5% CO_2_.

### Generation of stable two color reporter cell lines

The mNG2_1*−*10_ fragment was a gift from Siyu Feng ^47^ and was transfected using TransIT-LTI Transfection Reagent (Mirus) and packaged with pNTI673 and pNTI674 (Table S1) in a HEK 293T Lenti-X cell line to generate lentiviral particles. mNG2_1*−*10_ was then stably integrated into a polyclonal dCas9-KRAB CRISPRi K562 cell line ^52^ by lentiviral transduction. Dual color *CEBPA* reporter constructs (WT, ΔuORF start, ΔLong start, Short UTR, TableS1) were then stably integrated into this cell line by Cas9-mediated integration into the *AAVS1* locus by simultaneous nucleofection of plasmids containing either targeting sgRNAS (AAVS1-T2 and AAVS1-T2, Table S3) and spCas9^88^ and selected using 1 *µ*g/mL Puromycin (Invivogen) to generate stable integrants. All lines were subsequently monoclonally isolated.

### Generation of stable three color reporter cell lines

An IRFP-670 construct was stably integrated into the CRISPRi (mNG2_1*−*10_) cell line by Sleeping-beauty transposition ^89^ (Sleeping-beauty expression vector, Table S1), selected using 800 *µ*g/mL G418 (Invivogen) followed by monoclonal isolation. Generation of three color reporter lines was achieved by stable integration of dual color reporter constructs into the *AAVS1* locus in this cell line (as above).

### Flow cytometry

Cells were harvested for flow cytometry analysis by centrifugation (1500 RPM for 5 minutes at room temperature) followed by resuspension in PBS supplemented by 1% FBS and 1mM HEPES. All analyses were done on a LSR Fortessa Analyzer (BD Biosciences). Cells were initially gated on forward (FSC) and side scatter (SSC) (Figure S1A) and positive events were determined by a threshold based on negative (no stain) and single color control cells. Green, red and IRFP fluorescence was detected on the FITC (530/30 nM), PE-Texas Red (610/20 nM) and APC-Cy7 (780/60 nM) channels, respectively.

### FACS-based CRISPRi screen

CRISPRi sublibrary screens were performed using four, compact BFP-tagged CRISPRi sublibraries containing 5 sgRNAs per TSS (Addgene, Cat#83971-3 and #83975) expressed in the pCRISPRi-v2 expression vector (Addgene, Cat#84832). Plasmid sublibraries were separately packaged in HEK 293T Lenti-X cells and transduced into the CRISPRi dual color reporter line at an MOI *<* 1 where the percentage of transduced cells by BFP expression after 2 days post-transduction was 20%-30%. At 2 days post-transduction, we performed fluorescence activated cell sorting (FACS) using an Aria Fusion (BD Biosciences) to select for cells expressing BFP. Cells with the highest (*∼*20%) BFP expression were collected and recovered in RPMI 1640 for 6 days post-FACS. Approximately 10 million cells were collected per sublibrary, maintaining an average sgRNA coverage of at least 500 cells per sgRNA.

At 6 days post-BFP selection, cells were again sorted using a FACS Aria Fusion based on the ratio of green/red fluorescence from our *CEBPA* dual color reporter line. Approximately 40 million cells per sublibrary transduction were sorted into four, distinct green/red bins (*∼*20-25% of cells in each bin), with each bin containing *∼*8-10 million cells to ensure an average sgRNA/cell coverage of at least 500. Genomic DNA was immediately harvested from these cells using the DNeasy Blood and Tissue kit (Qiagen, 69504) and sgRNA fragments were isolated by SbfI (New England Biolabs) restriction digestion and Ampure bead size selection then amplified by PCR for deep sequencing as described in ^53^. The sgRNAs were sequenced on an Illumina HiSeq-4000 using custom primers.

### CRISPRi screen processing and data analysis

Sequencing reads were trimmed to remove adapter sequences using Cutadapt ^90^ and trimmed sgRNAs were counted using MAGeCK ^91,92^. Raw sgRNA counts were then used as input to DESeq2^93^ to calculate enrichment scores (Isoform Shift scores) in which each fluorescent bin (FR, NR, NG, FG) was represented as a numeric covariate in the linear model such that:

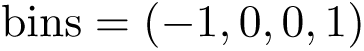

This assumes that sgRNA counts in the FR and FG bins (at the extremes of the fluorescent ratio distribution) have a constant multiplicative change with respect to the middle (NR and NG) bins.

Gene Ontology analysis was performed using PANTHER ^94–96^ with background lists representing the genes targeted by each hCRISPRa-v2 sublibrary.

### Individual validation of sgRNA-mediated phenotypes

Individual sgRNA expression vectors were cloned by first annealing complementary synthetic oligonucleotide sequences containing each sgRNA protospacer (Table S3) flanked by BstXI (New England Biolabs) and BlpI (New England Biolabs) restriction sites (Integrated DNA Technologies). Each double-stranded annealed pair was then ligated into a BstXI/BlpI-digested pCRISPRi-v2 expression vector containing a BFP cassette. Each sgRNA expression vector was then packaged into lentiviruses in HEK 293T Lenti-X cells and were individually transduced into either two or three color *CEBPA* CRISPRi reporter lines at a MOI *<* 1, resulting in *∼*20-30% infected cells by BFP expression. Cells were sorted on BFP 2 days post-transduction to select for sgRNA expression and allowed to recover in RPMI 1640 for 6 days. Reporter expression was then measured by flow cytometry.

### Western Blots

Cells were collected by centrifugation (1500 RPM for 5 minutes at room temperature), washed once with PBS, centrifuged again and lysed with buffer (140 mM KCl, 10 mM HEPES, 5 mM MgCl_2_, 1% TritonX-100, 1 mM TCEP, 2 U/ul Turbo DNAse) on ice for 30 minutes. Lysates were then clarified by centrifugation (14000 RPM for 10 minutes at 4*^◦^*C). Protein lysates were separated on Bolt 4%-8% Bis-Tris gels (Thermo Fisher Scientific) then transferred onto nitrocellulose membranes. Membranes were blocked with 5% milk in TBST (0.05% Tween-20) for 1 hour at room temperature. Primary antibodies were incubated overnight at 4*^◦^*C and secondary antibodies for 1 hour at room temperature. CEBPA protein was probed using a primary CEBPA antibody (1:1000, CST #2295), V5 epitope tags were probed by a primary V5-tag antibody (1:2000, CST #13202) and *β*-actin loading controls were probed by a primary, *β*-actin conjugated to HRP (1:2000, CST #12620). A HRP-conjugated anti-rabbit IgG (1:2000, CST #7074) was used as a secondary antibody against all primary antibodies. All blots were developed with SuperSignal West Dura Extended Duration Substrate (Thermo Scientific) and were visualized by a FluorChem R imaging system (ProteinSimple).

### PELO re-expression rescue assays

The HaloTag and PELO CDS was fused to the FKBP12 destabilizing domain (DD) to generate (N-terminal) DD-HaloTag and PELO-DD (C-terminal) constructs and were stably integrated into K562 CRISPRi dual color reporter lines by Sleeping-beauty transposition ^89^ (Sleeping-beauty expression vector, Table S1). Polyclonal cells were selected using 200 *µ*g/mL Hygromycin (Invivogen). Single sgRNAs were then packaged and transduced (as described above) into these cell lines and sorted for BFP-tagged sgRNA expression. Cells were then treated with 1 *µ*M Shield1 (Takara Bio Cat#632189) 5 days post-BFP sort then harvested for flow cytometry and Western blot analysis 3 days post-Shield1 treatment. Because of the incomplete destabilization of the PELO-DD construct (due to the necessity of placing the DD-tag at the C-terminus), we were unable to use a noninduced condition as a point of comparison (Figure S5A).

### Ribosome profiling and RNA-sequencing

For ribosome profiling and matched RNA sequencing, K562 CRISPRi dual color reporter cells were first transduced with either a nontargeting sgRNA or the top scoring *PELO* sgRNA (see individual sgRNA knockdown validation) in triplicate and were grown in T150 flasks (Corning) for 6 days post-BFP selection. Cells (5.0*x*10^6^ nontargeting sgRNA and 2.5*x*10^6^ PELO sgRNA per replicate) were harvested as previously described ^72^ without addition of cycloheximide and a sample of the lysate was taken for RNA sequencing. Ribosome profiling was done as described in ^72^.

For matched RNA sequencing, cells were harvested by phenol-chloroform extraction and processed according to the NEB Ultra II Directional RNA Sample Prep Kit (New England Biolabs, #E7760S).

The ribosome profiling and total RNA sequencing samples were sequenced on an Illumina NovaSeq instrument.

### Ribosome profiling sequencing analysis

Ribosome profiling reads were first trimmed using Cutadapt and aligned to ribosomal RNA (rRNA) and transfer RNA (tRNA) references using Bowtie2^97^. The remaining reads were subsequently aligned to the transcriptome using STAR ^98^. Transcriptome-based alignments were then filtered to exclude the first 15 and last 5 codons due to the accumulation of initiating and terminating ribosomes. Finally, footprint abundance was quantified from these alignments by RSEM ^99^. Differential expression and translation efficiency analysis was conducted using DESeq2^93^.

---

**Supplementary Figure 1:**
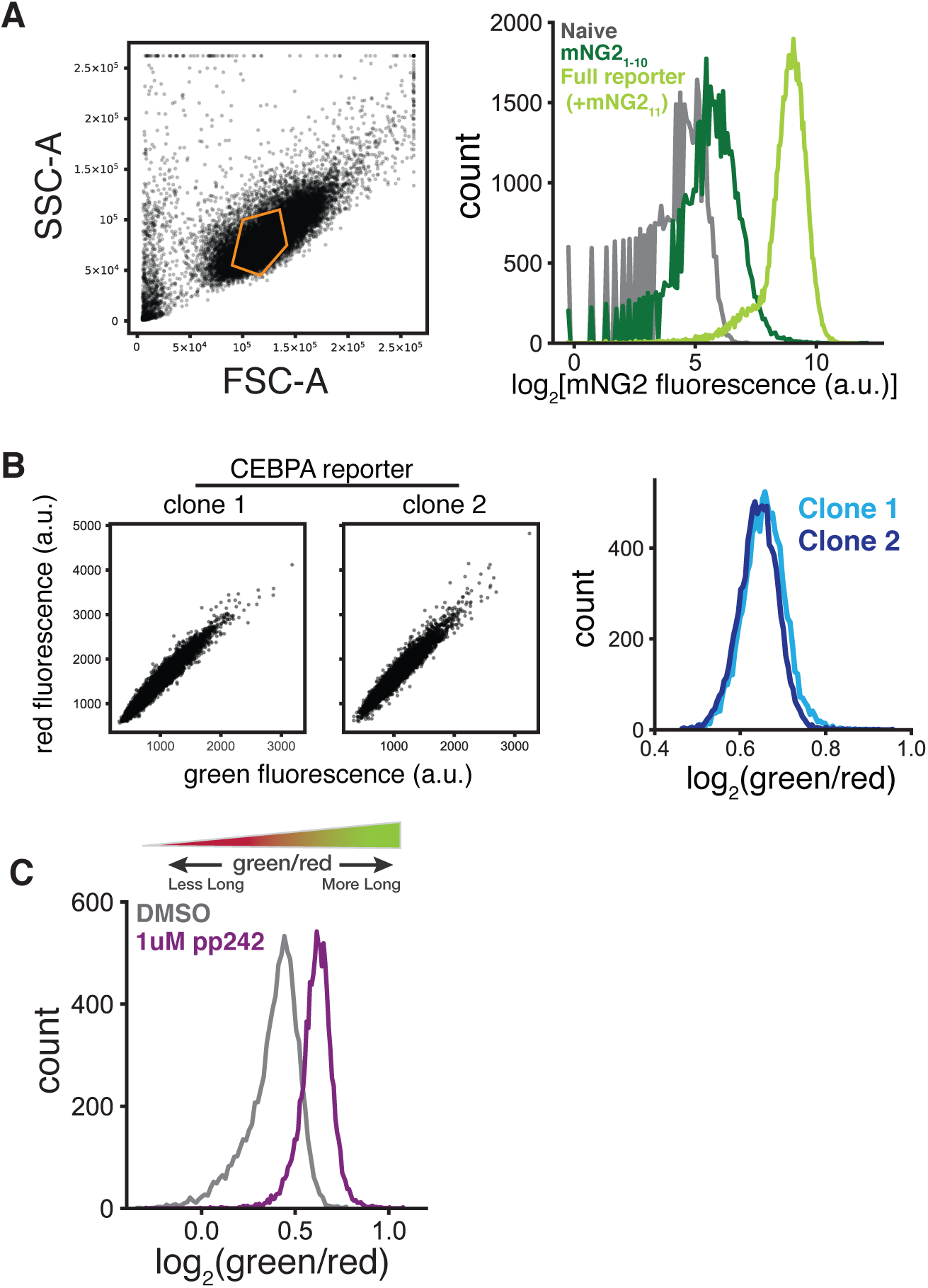
Validation of *CEBPA* two color reporter. (A) Forward (FSC-A) and side scatter (SSC-A) gating scheme for all flow cytometry measurements (left). Distribution of mNeonGreen2 (mNG2) fluorescence by flow cytometry from cell lines containing neither mNG2 fragment (naive), a cell line constitutively expressing only the mNG2_1*−*10_ fragment or a cell line expressing both mNG2_1*−*10_ and the *CEBPA* reporter containing the mNG2_11_ fragment (Full reporter) (right). (B) Scatterplots of green and red fluorescence from two clonal, stable *CEBPA* two color reporter cell lines (left) and green/red fluorescence distributions from these two clonal lines by flow cytometry (right). (C) Green/red fluorescence distributions in wild type two color reporter cell lines treated with either DMSO or 1 *µ*M PP242 (Sigma-Aldrich). Cells were harvested and assayed by flow cytometry 24h post-treatment.

**Supplementary Figure 2:**
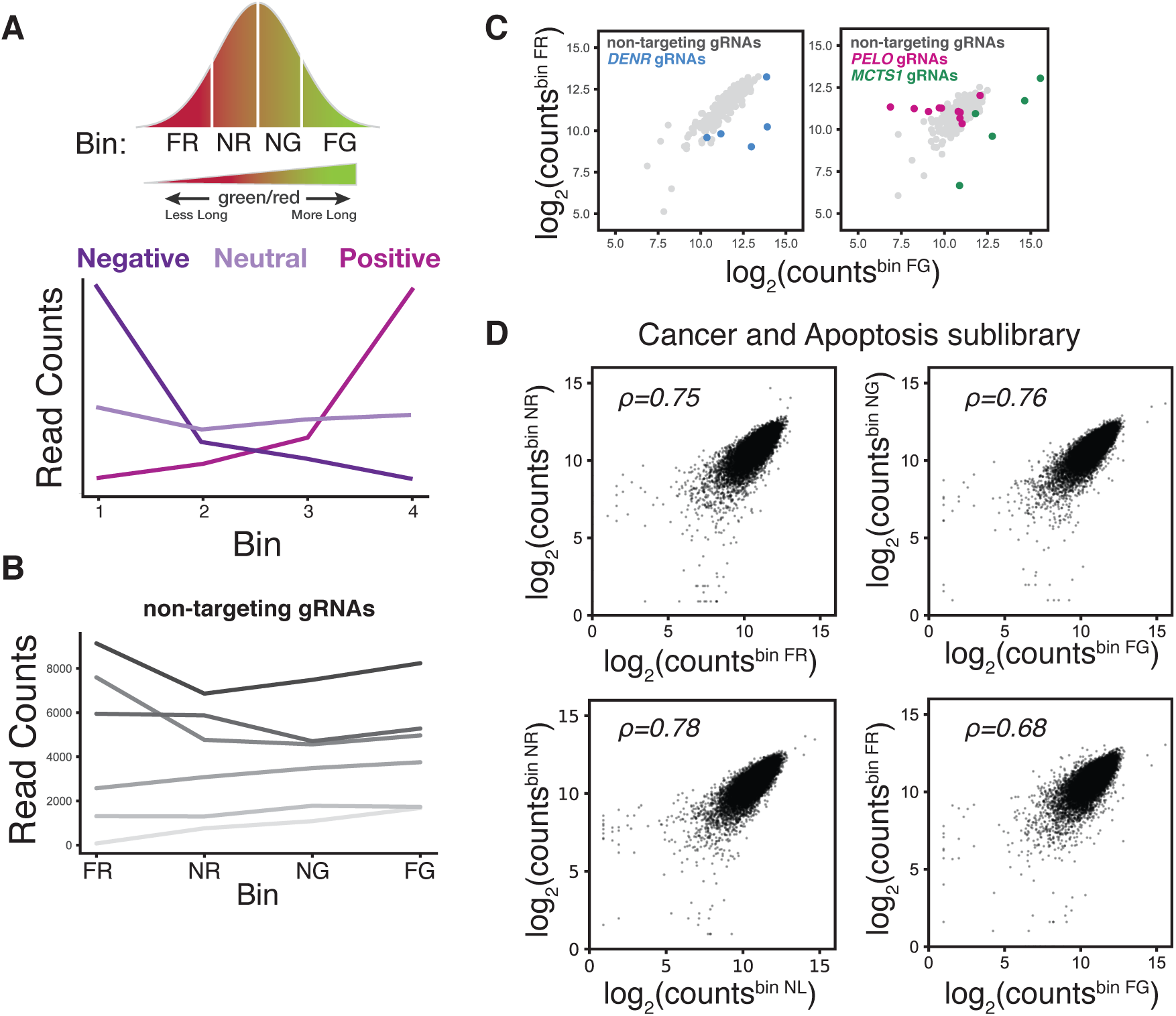
Validation of FACS-based CRISPRi sublibrary screens. (A) Schematic representation of possible, functional outcomes of sgRNA-mediated knockdown on the two color reporter. Bin labels reflect green/red distribution. FR: far red; NR: near red; NG: near green; FG: far green. sgRNAs that are enriched in cells with higher green/red ratios (NG and FG) may be changing the ratio by shifting expression towards the long isoform. We consider this class of sgRNAs as positive regulators of short isoform expression. In contrast, sgRNAs that are enriched in cells with lower green/red ratios (NR and FR) may be changing the ratio by shifting isoform expression towards the short isoform. We consider this class of sgRNAs as negative regulators of short isoform expression. sgRNAs that are uniformly distributed across all bins either have no effect on isoform usage or affect both isoforms equivalently. We classify these sgRNAs as being neutral. (B) sgRNA read count distribution across all bins for 6 nontargeting sgRNAs from the Gene Expression sublibrary. (C) Comparison of sgRNA read counts between the Far Red (FR) and Far Green (FG) bins in the Gene Expression sublibrary (left) and Cancer and Apoptosis sublibrary (right). Individual sgRNAs against *DENR*, *MCTS1* and *PELO* are highlighted. (D) Comparison of sgRNA read counts between indicated fluorescent bins in the Cancer and Apoptosis sublibrary (as in Figure 2D). Each point represents one distinct sgRNA.

**Supplementary Figure 3:**
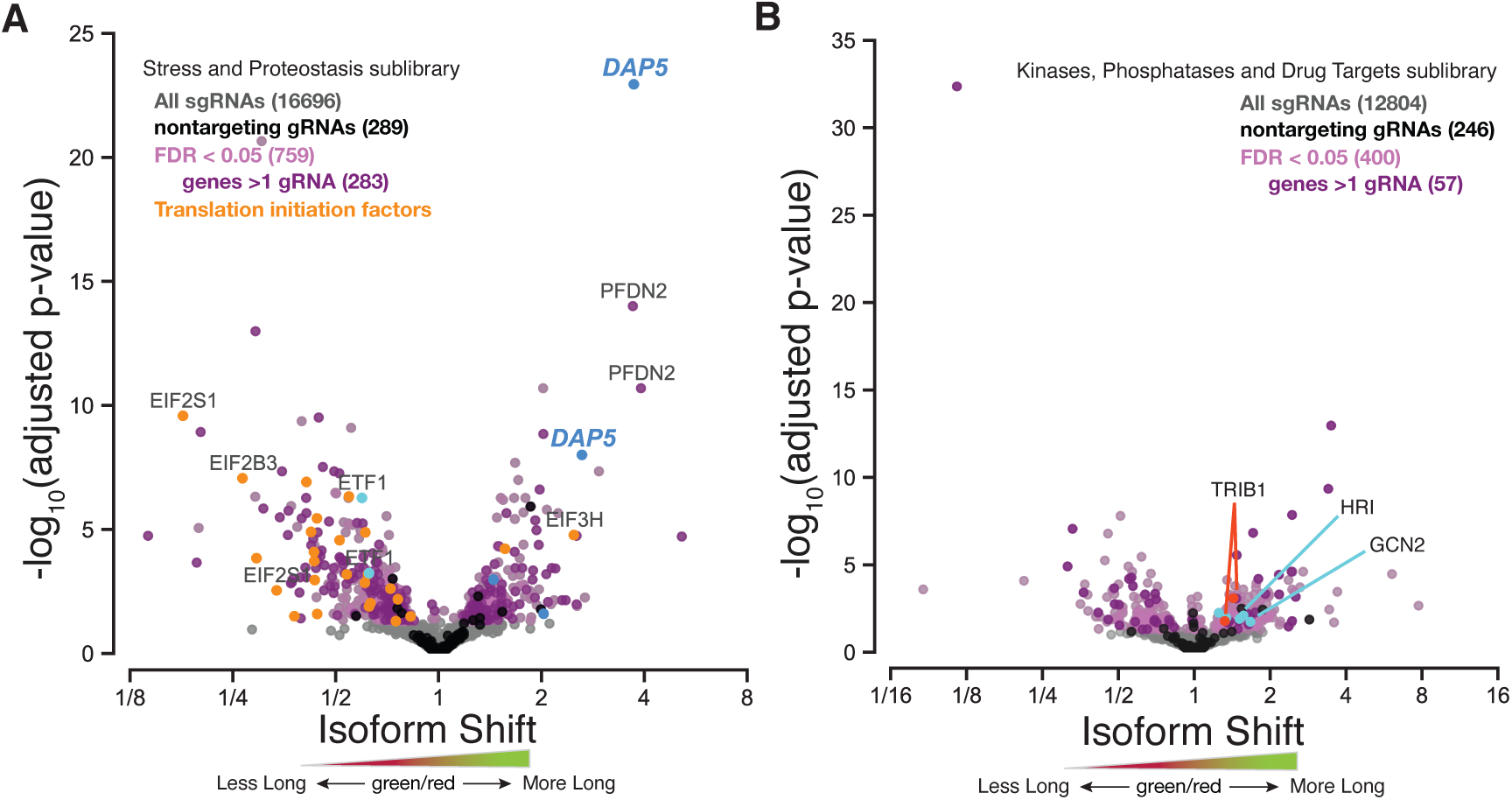
Additional FACS-based CRISPRi sublibrary screen results. (A) Stress and Proteostasis sgRNA sublibrary profile representing the relative shift in long and short isoform usage. Each point represents a single sgRNA with sgRNAs against *eIF4G2/DAP5*, translation termination factor 1 (*ETF1*) and translation initiation factors highlighted. Colors indicate cutoffs for significance (false discovery rate, FDR *<* 0.05), genes with at least 2 sgRNAs and nontargeting sgRNAs. (B) Kinases, Phosphatases and Drug Targets sgRNA sublibrary profile representing the relative shift in long and short isoform usage, as in (A). sgRNAs against the *eIF2α* kinases, *HRI* and *GCN2*, and the pseudokinase *TRIB1* are indicated.

**Supplementary Figure 4:**
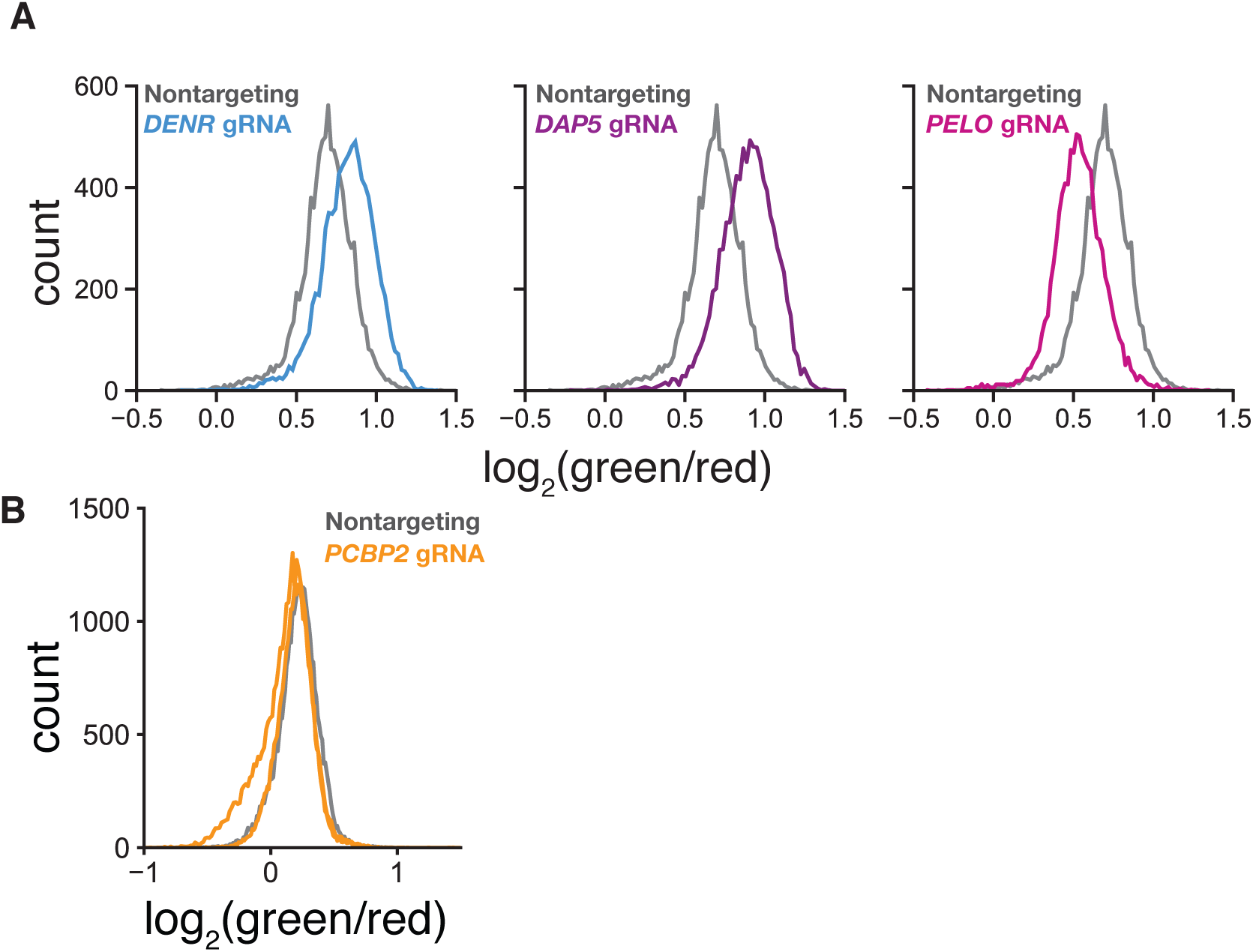
Validation of individual sgRNA mediated knockdowns in two color reporter cell lines. (A) Distribution of green/red fluorescence of a second, clonal two color reporter cell line transduced with the top scoring sgRNAs against either *DENR*, *DAP5* or *PELO*. (B) Distribution of green/red fluorescence of the two color reporter cell line transduced with 2 sgRNAs against *PCBP2*.

**Supplementary Figure 5:**
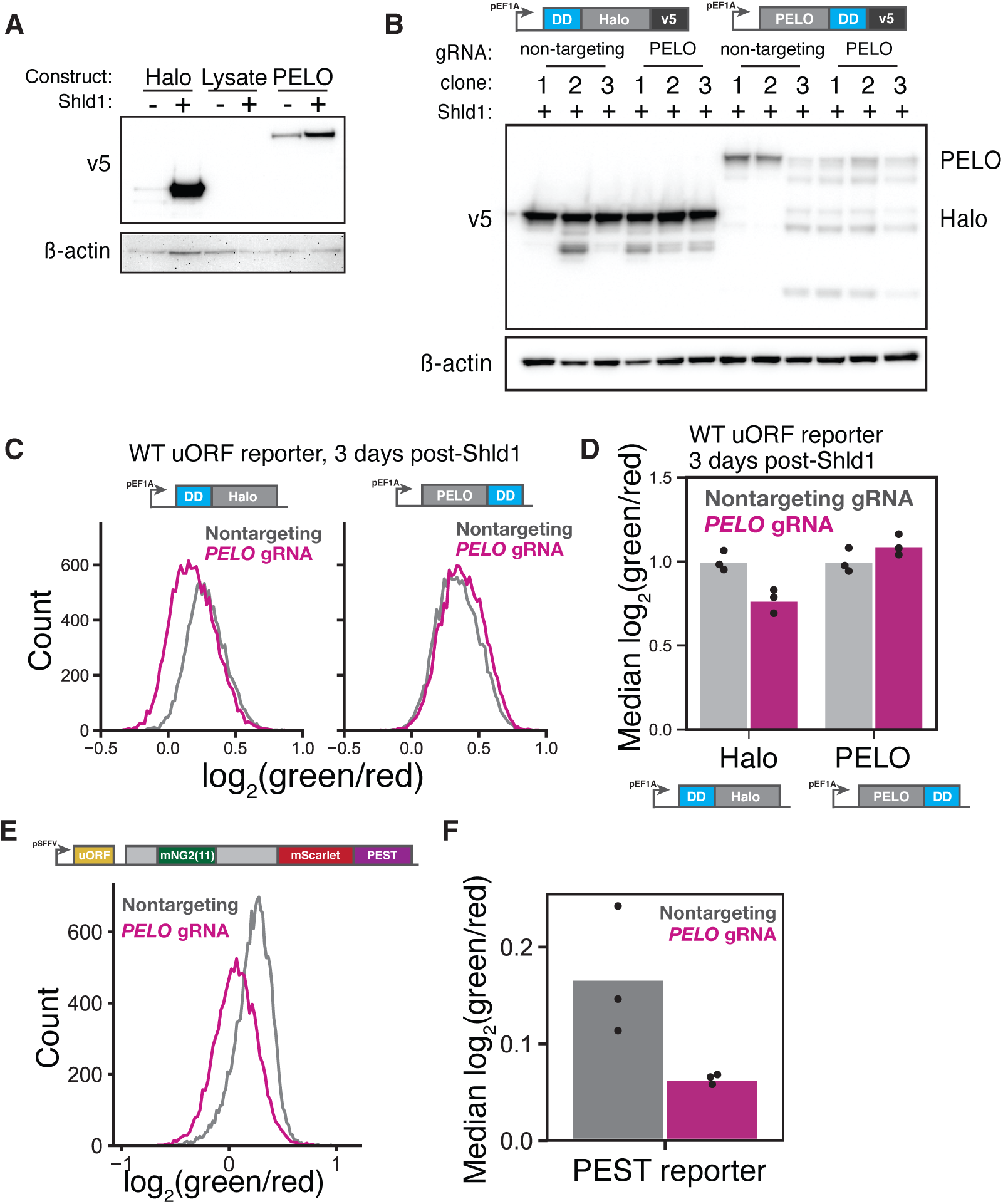
*PELO* knockdown phenotype does not depend on uORF start codon context or differential protein stability. (A) Western blot of naive K562 cells nucleofected with V5-tagged constructs containing either HaloTag or PELO fused to the FKBP12 destabilizing domain (DD) relative to *β*-actin loading control. Cells were either left untreated or treated with 1 *µ*M Shield1 (Shld1) (Takara Bio) then harvested 72h post-treatment for Western blot. (B) Western blot of two color reporter cell lines stably expressing either a V5-tagged DD-HaloTag or PELO-DD fusion transduced with either a nontargeting sgRNA or a *PELO* sgRNA. *β*-actin was used as a loading control. DD-fusion cell lines were first transduced with sgRNAs, allowed to recover for 5 days then treated with 1 *µ*M Shld1. Cells were harvested for Western blot 72h post-Shld1 treatment. Clones represent separate, independent transductions. (C) Distribution of green/red fluorescence by flow cytometry in a two color reporter cell line harboring the native, uORF Kozak sequence (ctcgccATGc) stably expressing either a DD-HaloTag or PELO-DD fusion construct and transduced with either a nontargeting or a *PELO* sgRNA. Cells were transduced and treated as in (B). (D) Median green/red fluorescence measurements of the cell lines in (C), n = 3. (E) Distribution of green/red fluorescence by flow cytometry in a two color reporter cell line bearing a C-terminal PEST sequence transduced with either a nontargeting or *PELO* sgRNA. (F) Median green/red fluorescence measurements of PEST-fusion two color reporters in (E), n = 3.

**Supplementary Figure 6:**
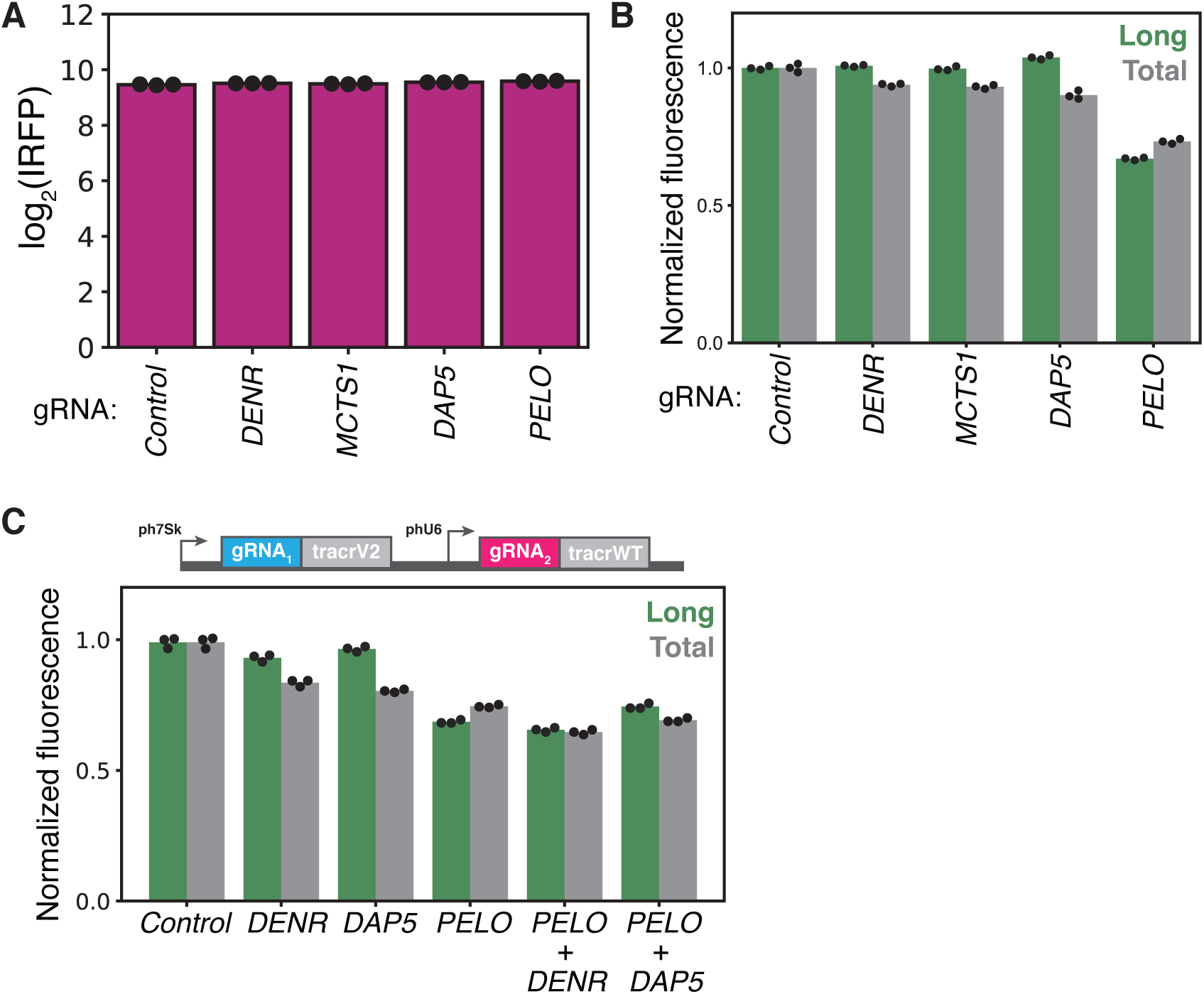
Validation of *CEBPA* three color reporter. (A) Flow cytometry measurements of median IRFP670 fluorescence in stable cell lines expressing the three color reporter transduced with indicated sgRNAs, n = 3. (B) Flow cytometry measurements of median green and red fluorescence normalized to IRFP670 in stable cell lines expressing the wild type three color reporter transduced with indicated sgRNAs, n = 3, as in Figure 5B. Fluorescent values were normalized to the median values in the negative, nontargeting control. (C) Flow cytometry measurements of median green and red fluorescence normalized to IRFP in stable cell lines expressing the wild type three color reporter transduced with indicated dual sgRNAs, n = 3, as in Figure 5C. Fluorescent measurements were normalized as in (B).

**Supplementary Figure 7:**
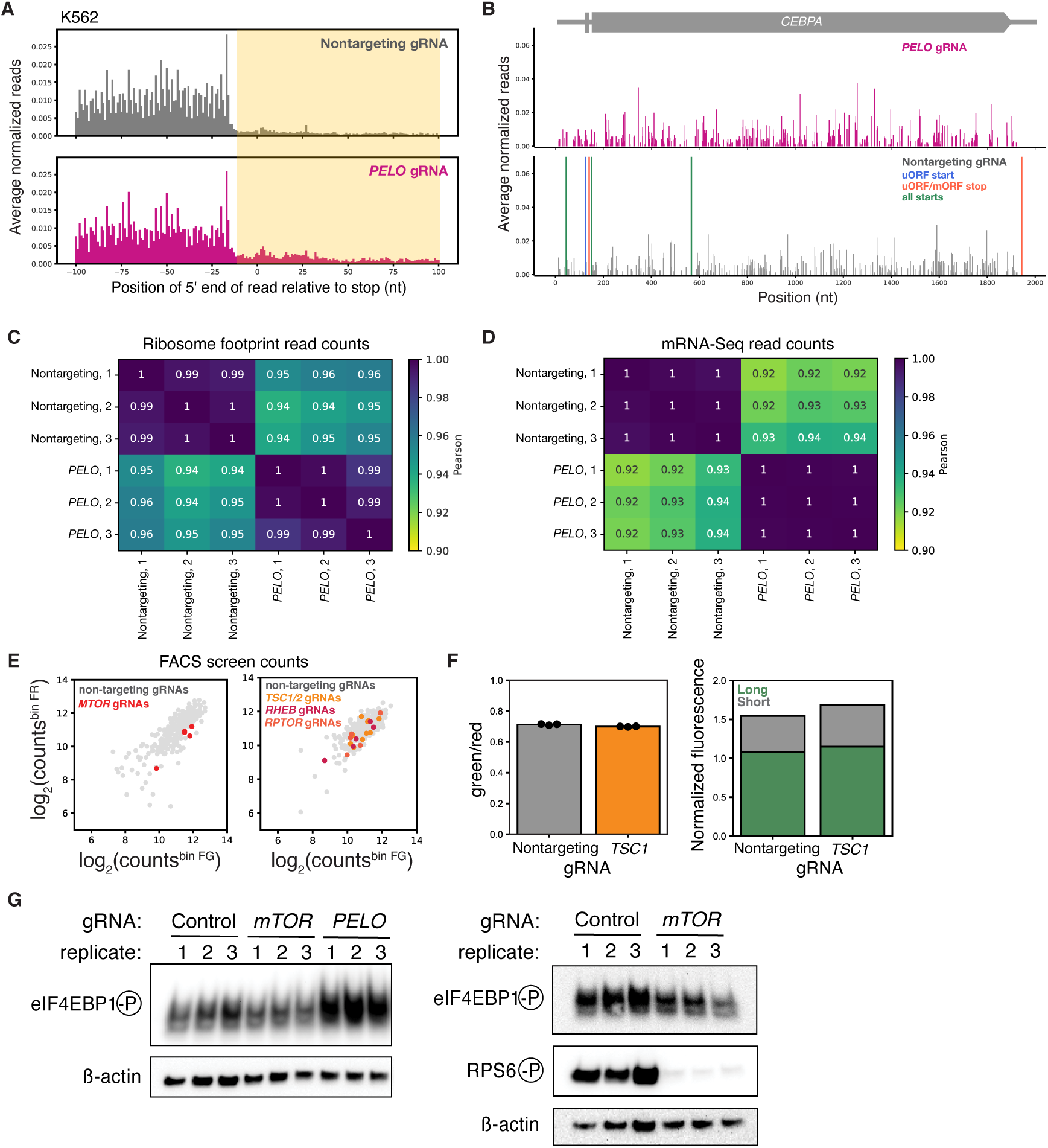
*PELO* knockdown triggers mTOR activation. (A) Metagene analyses of ribosome occupancy in either the nontargeting control or *PELO* knockdown in two color reporter line. Positions indicate the 5’ end of footprints relative to stop codons. Read counts were normalized by the total ribosome footprint count in each library and the median count was determined across replicates, n=3. (B) Ribosome footprint profile of the *CEBPA* reporter in cells expressing either a nontargeting sgRNA or a *PELO* sgRNA. Start and stop codons are indicated and counts were normalized as in (A). (C) Pearson correlation of ribosome footprint counts between indicated libraries. (D) Pearson correlation of ribosome profiling matched mRNA-seq counts between indicated libraries. (E) Comparison of sgRNA read counts between the Far Red (FR) and Far Green (FG) bins in the Kinases, Phosphatases and Drug Targets sublibrary (left) and Cancer and Apoptosis sublibrary (right). Individual sgRNAs against *MTOR*, *RHEB*, *TSC1*, *TSC2* and *RAPTOR* are highlighted. (F) Flow cytometry measurement of three color reporter cell lines transduced with either a sgRNA against *TSC1* or a nontargeting sgRNA. Median green/red ratio is displayed, n = 3 (left). Flow cytometry measurements of median IRFP670 fluorescence in stable cell lines expressing the three color reporter transduced with indicated sgRNAs, n = 3 (right). (G) Western blot of three color reporter cell lines transduced with the indicated dual sgRNAs as in Figure 6D. *β*-actin was used as a loading control. Replicates represent separate, independent transductions.

## References

[1] Marilyn Kozak. Point mutations define a sequence flanking the aug initiator codon that modulates translation by eukaryotic ribosomes. Cell, 44:283–292, 1986.

[2] Alan G Hinnebusch. Molecular mechanism of scanning and start codon selection in eukaryotes. Microbiol. Mol. Biol. Rev., 75:434–467, 2011.

[3] Alan G Hinnebusch, Ivaylo P Ivanov, and Nahum Sonenberg. Translational control by 5’-untranslated regions of eukaryotic mRNAs. Science, 352:1413–1416, 2016.

[4] Alex V Kochetov, Akinori Sarai, Igor B Rogozin, Vladimir K Shumny, and Nikolay A Kolchanov. The role of alternative translation start sites in the generation of human protein diversity. Mol Gen Genomics, 273:273–291, 2005.

[5] Mari L Shinohara, Hye-Jung Kim, June-Ho Kim, Virgilio A Garcia, and Harvey Cantor. Alternative translation of osteopontin generates intracellular and secreted isoforms that mediate distinct biological activities in dendritic cells. Proc. Natl. Acad. Sci., 105:7235–7239, 2008.

[6] Nicholas T Ingolia, Liana F Lareau, and Jonathan S Weissman. Ribosome profiling of mouse embryonic stem cells reveals the complexity and dynamics of mammalian proteomes. Cell, 147:789–802, 2011.

[7] Gloria A Brar, Moran Yassour, Nir Friedman, Aviv Regev, Nicholas T Ingolia, and Jonathan S Weissman. High-resolution view of the yeast meiotic program revealed by ribosome profling. Science, 335:552–557, 2011.

[8] Christoph Burkart, Jun-Bao Fan, and Dong-Er Zhang. Two independent mechanisms promote expression of an n-terminal truncated USP18 isoform with higher delSGylation activity in the nucleus. J Biol Chem, 287:4883–4893, 2012.

[9] Luis J Cocka and Paul Bates. Identification of alternatively translated tetherin isoforms with differing antiviral and signaling activities. PLoS Pathog., 8, 2012.

[10] Nicholas T Ingolia, Gloria A Brar, Noam Stern-Ginossar, Michael S Harris, Gäelle JS Talhouarne, Sarah E Jackson, Mark R Wills, and Jonathan S Weissman. Ribosome profiling reveals pervasive translation outside of annotated protein-coding genes. Cell Rep, 8:1365– 1379, 2014.

[11] Alexander P Fields, Edwin H Rodriguez, Marko Jovanovic, Noam Stern-Ginossar, Brian J Haas, Philipp Mertins, Raktima Raychowdhury, Nir Hacohen, Steven A Carr, Nicholas T Ingolia, Aviv Regev, and Jonathan S Weissman. A regression-based analysis of ribosome-profiling data reveals a conserved complexity to mammalian translation. Mol. Cell, 60:816–827, 2015.

[12] Jin Chen, Andreas-David Brunner, J Zachery Cogan, James K Nunez, Alexander P Fields, Britt Adamson, Daniel N Itzhak, Jason Y Li, Mattias Mann, Manuel D Leonetti, and Jonathan S Weissman. Pervasive functional translation of noncanonical human open reading frames. Science, 367:1140–1146, 2020.

[13] Patric Descombes and Ueli Schibler. A liver-enriched transcriptional activator protein, LAP, and a trascriptional inhibitory protein, LIP, are translated from the same mrna. Cell, 67: 569–579, 1991.

[14] Cor F Calkhoven, Christine Müller, and Achim Leutz. Translational control of *C/EBPA* and *C/EBPB* isoform expression. Genes Dev, 25:1920–1932, 2000.

[15] Bo Li and Colin N Dewey. p53 stability and activity is regulated by mdm2-mediated induction of alternative p53 translation products. Nat Cell Biol., 4:462–467, 2002.

[16] Nick Z Lu and John A Cidlowski. Translational regulatory mechanisms generate n-terminal glucocorticoid receptor isoforms with unique transcriptional target genes. Mol. Cell, 5:331–342, 2005.

[17] Mary Jane Tsang and Iain M Cheeseman. Alternative Cdc20 translational isoforms bypass the spindle assembly checkpoint to control mitotic arrest duration. BioarXiv, 2021.

[18] Sky W Brubaker, Anna E Gauthier, Eric W Mill, Nicholas T Ingolia, and Jonathan C Kagan. A bicistronic *MAVS* transcript highlights a class of truncated variants in antiviral immunity. Cell, 156:800–811, 2014.

[19] Sarah E Calvo, David J Pagliarini, and Vamsi K Mootha. Upstream open reading frames cause widespread reduction of protein expression and are polymorphic among humans. Proc. Natl. Acad. Sci., 106:7507–7512, 2009.

[20] Patrick McGillivray, Russell Ault, Mayur Pawashe, Robert Kitchen, Suganthi Balasubramanian, and Mark Gerstein. A comprehensive catalog of predicted functional upstream open reading frames in humans. Nucleic Acids Res, 46:3326–3338, 2018.

[21] David R Morris and Adam P Geballe. Upstream open reading frames as regulators of mrna translation. Molecular and cellular biology, 20(23):8635–8642, 2000.

[22] Klaus Wethmar. The regulatory potential of upstream open reading frames in eukaryotic gene expression. Wiley Interdisciplinary Reviews: RNA, 5(6):765–768, 2014.

[23] Timothy G Johnstone, Ariel A Bazzini, and Antonio J Giraldez. Upstream ORFs are prevalent translational repressors in vertebrates. The EMBO Journal, 35:706–723, 2018.

[24] Jianhong Cao and Adam P Geballe. Coding sequence-dependent ribosomal arrest at termination of translation. Molecular and Cellular Biology, 16(2):603–608, 1996.

[25] Zhong Wang, Anthony Gaba, and Matthew S Sachs. A highly conserved mechanism of regulated ribosome stalling mediated by fungal arginine attenuator peptides that appears independent of the charging status of arginyl-trnas. Journal of Biological Chemistry, 274(53): 37565–37574, 1999.

[26] Sara K Young and Ronald C Wek. Upstream open reading frames differentially regulate gene-specific translation in the integrated stress response. Journal of Biological Chemistry, 291(33): 16927–16935, 2016.

[27] Ivaylo P Ivanov, Byung-Sik Shin, Gary Loughran, Ioanna Tzani, Sara K Young-Baird, Chune Cao, John F Atkins, and Thomas E Dever. Polyamine control of translation elongation regulates start site selection on antizyme inhibitor mrna via ribosome queuing. Mol. Cell, 70: 254–264, 2018.

[28] Yizhu Lin, Gemma E May, Hunter Kready, Lauren Nazzaro, Mao Mao, Pieter Spealman, Yehuda Creeger, and C Joel McManus. Impacts of uorf codon identity and position on translation regulation. Nucleic acids research, 47(17):9358–9367, 2019.

[29] Ty A Bottorff, Heungwon Park, Adam P Geballe, and Arvind Rasi Subramaniam. Translational buffering by ribosome stalling in upstream open reading frames. PLoS Genetics, 18(10): e1010460, 2022.

[30] Krishna M Vattem and Ronald C Wek. Reinitiation involving upstream ORFs regulates *ATF4* mRNA translation in mammalian cells. Proc. Natl. Acad. Sci., 101:11269–11274, 2004.

[31] Phoebe D Lu, Heather P Harding, and David Ron. Translation reintiation at alternative open reading frames regulates gene expression in an integrated stress response. J. Cell Biol., 167: 27–33, 2004.

[32] Thomas Pabst, Beatrice U Mueller, Pu Zhang, Hanna S Radomska, Sailaja Narravula, Susanne Schnittger, Gerhard Behre, Wolfgang Hiddemann, and Daniel G Tenen. Dominant-negative mutations of *CEBPA*, encoding CCAAT/enhancer binding protein-*α*(C/EBP*α*), in acute myeloid leukemia. Nat Genet, 27:263–270, 2001.

[33] Anna Mae Diehl, Philip Michaelson, and Shi Qi Yang. Selective induction of ccaat/enhancer binding protein isoforms occurs during rat liver development. Gastroenterology, 106:1625–1637, 1994.

[34] Rana Basabi, Yuhong Xie, David Mischoulon, Nancy L R Bucher, and Stephen R Farmer. The dna binding activity of C/EBP transcription factors is regulated in the g1 phase of the hepatocyte cell cycle. Gastroenterology, 270:18123–18132, 1995.

[35] Peggy Kirstetter, Mikkel B Schuster, Oksana Bereshchenko, Susan Moore, Heidi Dvinge, Elke Kurz, Kim Theilgaard-Mönch, Månsson, Thomas Pedersen, Thomas Pabst, Evelin Schrock, Bo T Porse, Sten Eirik W Jacobsen, Paul Bertone, Daniel G Tenen, and Claus Nerlov. Modeling of c/ebp*α* mutant acute myeloid leukemia reveals a common expression signature of committed myeloid leukemia-initiating cells. Cancer Cell, 17:299–310, 2008.

[36] Janus S Jakobsen, Linea G Laursen, Mikkel B Schuster, Sachin Pundhir, Erwin Schoof, Ying Ge, Teresa D’altri, Kristoffer Vitting-Seerup, Nicholas Rapin, Coline Gentil, Johan Jendholm, Kim Theilgaard-Mönch, Kristian Reckzeh, Lars Bullinger, Konstanze Döhner, Peter Hokland, Jude Fitzgibbon, and Bo T Porse. Mutant CEBPA directly drives the expression of the targetable tumor-promoting factor CD73 in AML. Sci. Adv., 5, 2019.

[37] A Fasan, C Haferlach, T Alpermann, S Jeromin, V Grossman, C Eder, S Weissman, F Dicker, A Kohlmann, S Schindela, W Kern, T Haferlach, and S Schnittger. The role of different genetic subtypes of CEBPA mutated AML. Leukemia, 28:794–803, 2014.

[38] Sibylle Schleich, Katrin Strassburger, Philipp C Janiesch, Tatyana Koledachkina, Katharine K Miller, Katharina Haneke, Yong-Sheng Cheng, Katrin Küchler, George Stoecklin, Kent E Duncan, and Aurelio A Teleman. DENR-MCT-1 promotes translation re-initiaiton downstream of uorfs to control tissue growth. Nature, 512:208–212, 2014.

[39] Jonathan Bohlen, Liza Harbrecht, Saioa Blanco, Katharina Clemm von Hohenberg, Kai Fenzl, Günter Kramer, Bernd Bukau, and Aurelio A Teleman. *DENR* promotes translation reinitiation via ribosome recycling to drive expression of oncogenes including *ATF4*. Nature Comm, 11, 2020.

[40] Deepika Vasudevan, Sarah D Neuman, Amy Yang, Lea Lough, Brian Brown, Arash Bashirullah, Timothy Cardozo, and Hyung Don Ryoo. Translational induction of ATF4 during integrated stress response requires noncanonical initiation factors eIF2D and DENR. Nature Comm, 11, 2020.

[41] Vera P Pisareva, Maxim A Skabkin, Christopher U T Hellen, Tatyana V Pestova, and Andrey V Pisarev. Dissociation by Pelota, Hbs1 and ABCE1 of mammalian vacant 80s ribosomes and stalled elongation complexes. The EMBO Journal, 30:1804–1817, 2011.

[42] Thomas Becker, Sibylle Franckenberg, Stephan Wickles, Christopher J Shoemaker, Andreas M Anger, Jean-Paul Armache, Heidemarie Sieber, Charlotte Ungewickell, Otto Berninghausen, Ingo Daberkow, Annette Karcher, Michael Thomm, Karl-Peter Hopfner, Rachel Green, and Roland Beckman. Structural basis of highly conserved ribosome recycling in eukaryotes and archaea. Nature, 482:501–506, 2012.

[43] Nicholas R Guydosh and Rachel Green. Dom34 rescues ribosomes in 3’ untranslated regions. Cell, 156:950–962, 2014.

[44] Eric W Mills, Jamie Wangen, Rachel Green, and Nicholas T Ingolia. Dynamic regulation of a ribosome rescue pathway in erythroid cells and platelets. Cell Rep, 17:1–10, 2016.

[45] Daphne S Bindels, Lindsay Haarbosch, Laura van Weeren, Marten Postma, Katrin E Wiese, Marieke Mastop, Sylvain Aumonier, Guillaume Gotthard, Antoine Royant, Mark A Hink, and Theodorus W J Gadella Jr. mScarlet: a bright monomeric red fluorescent protein for cellular imaging. Nat Methods, 14:53–56, 2017.

[46] Nathan C Shaner, Gerard G Lambert, Andrew Chammas, Yuhui Ni, Paula J Cranfill, Michelle A Baird, Brittney R Sell, John R Allen, Richard N Day, Maria Israelsson, Michael W Davidson, and Jiwu Wang. A bright monomeric green fluorescent protein derived from *Branchiostoma lanceolatum*. Nat Methods, 10:407–409, 2013.

[47] Siyu Feng, Sayaka Sekine, Veronica Pessino, Han Li, Manuel D. Leonetti, and Bo Huang. Improved split fluorescent proteins for endogenous protein labeling. Nature Comm, 8, 2017.

[48] Morris E Feldman, Beth Apsel, Aino Uotila, Robbie Lowith, Zachary A Knight, Davide Ruggero, and Kevan M Shokat. Active-site inhibitors of mtor target rapamycin-resistant outputs of mtorc1 and mtorc2. PLoS Biol, 7:e1000038, 2009.

[49] Carson C Thoreen, Seong A Kang, Jae Won Chang, Qingsong Liu, Jianming Zhang, Yi Gao, Laurie J Reichling, Taebo Sim, David M Sabatini, and Nathanael S Gray. An atp-competitive mammalian target of rapamycin inhibitor reveals rapamycin-resistant functions of mtorc1. J Biol Chem, 284:8023–8032, 2009.

[50] Justin B Kinney, Anand Murugan, Curtis G Callan Jr, and Edward C Cox. Using deep sequencing to characterize the biophysical mechanism of a transcriptional regulatory sequence. Proc. Natl. Acad. Sci., 107:9158–9163, 2010.

[51] William L Noderer, Ross J Flockhart, Aparna Bhaduri, Alexander J Diaz de Arce, Jiajing Zhang, Paul A Khavari, and Clifford L Wang. Quantitative analysis of mammalian translation initiation sites by FACS-seq. Mol Syst Biol., 10, 2014.

[52] Luke A Gilbert, Max A Horlbeck, Britt Adamson, Jacqueline E Villalta, Yuwen Chen, Evan H Whitehead, Carla Guimaraes, Barbara Panning, Hidde L Ploegh, Michael C Bassik, Lei S Qi, Martin Kampmann, and Jonathan S Weissman. Genome-scale crispr-mediated control for gene repression and activation. Cell, 159:647–661, 2014.

[53] Max A Horlbeck, Luke A Gilbert, Jacqueline E Villalta, Britt Adamson, Ryan A Pak, Yuwen Chen, Alexander P Fields, Chong Yon Park, Jacob E Corn, Martin Kampmann, and Jonathan S Weissman. Compact and highly active next-generation libraries for crispr-mediated gene repression and activation. eLife, 5:e19760, 2016.

[54] Melanie Weisser, Tanja Schäfer, Marc Leibundgut, Daniel Böhringer, Christopher Herbert Stanley Aylett, and Nenad Ban. Structural and functional insights into human reinitiation complexes. Mol. Cell, 67:447–456, 2017.

[55] Ivan B Lomakin, Elena A Stolboushkina, Anand T Vaidya, Chenguang Zhao, Maria B Garber, Sergey E Dmitriev, and Thomas A Steitz. Crystal structure of the human ribosome in complex with DENR-MCT-1. Cell Rep, 20:521–528, 2017.

[56] Yasar Luqman Ahmed, Sibylle Schleich, Jonathan Bohlen, Nicolas Mandel, Bernd Simon, Irmgard Sinning, and Aurelio A Teleman. DENR-MCTS1 heterodimerization and tRNA recruitment are required for translation reinitiation. PLoS Biol, 10, 2018.

[57] Ivan B Lomakin, Sergey E Dmitriev, and Thomas A Steitz. Crystal structure of the DENR-MCT-1 complex revealed zinc-binding site essential for heterodimer formation. Proc. Natl. Acad. Sci., 116:528–533, 2019.

[58] Sibylle Schleich, Julieta M Acevedo, von Katharina Clemm Hohenberg, and Aurelio A Teleman. Identification of transcripts with short stuorfs as targets for DENR-MCTS1-dependent translation in human cells. Sci Rep, 7, 2017.

[59] Ramona Weber, Leon Kleemann, Insa Hirschberg, Min-Yi Chung, Eugene Valkov, and Cátia Igreja. *DAP5* enables translation re-initiation on mRNAs with structured and uORF-containing 5’ leaders. Nature Comm, 13, 2022.

[60] Maya David, Tsviya Olender, Orel Mizrahi, Shira Weingarten-Gabbay, Gilgi Friedlander, Sara Meril, Nadav Goldber, Alon Savidor, Yishai Levin, Vered Salomon, Noam Stern-Ginossar, Shani Bialik, and Adi Kimchi. *DAP5* drives translation of specific mrna targets with upstream orfs in human embryonic stem cells. RNA, 2022.

[61] K Doma, Meenakshi and Roy Parker. Endonucleolytic cleavage of eukaryotic mRNAs with stalls in translation elongation. Nature, 440:561–564, 2006.

[62] Christopher J Shoemaker and Rachel Green. Kinetic analysis reveals the ordered coupling of translation termination and ribosome recycling in yeast. Proc. Natl. Acad. Sci., 108:E1392–E1398, 2011.

[63] Marc Graille, Maxime Chaillet, and Herman van Tilbeurgh. Structure of yeast dom34: a protein related to translation termination factor erf1 and involved in no-go decay. J Biol Chem, 293:7145–7154, 2008.

[64] Maxim A Skabkin, Olga V Skabkina, Vidya Dhote, Anton A Komar, Christoper UT Hellen, and Tatyana V Pestova. Activities of Ligatin and MCT-1/DENR in eukaryotic translation initiation and ribosomal recycling. Genes Dev, 24:1787–1801, 2010.

[65] David J Young, Desislava S Makeeva, Fan Zhang, Aleksandra S Anisimova, Elena A Stolboushkina, Fardin Ghobakhlou, Ivan N Shatsky, Sergey E Dmitriev, Alan G Hinnebusch, and Nicholas R Guydosh. Tma64/eIF2D,Tma20/MCT-1, and Tma22/DENR recycle post-termination 40s subunits *In Vivo*. Mol. Cell, 71:761–774, 2018.

[66] Laura A Banaszynski, Ling-Chun Chen, Lystranne A Maynard-Smith, A G Lisa Ooi, and Thomas J Wandless. A rapid, reversible, and tunable method to regulate protein function in living cells using synthetic small molecules. Cell, 126:995–1004, 2006.

[67] Jonathan Bohlen, Kai Fenzl, Günter Kramer, Bernd Bukau, and Aurelio A Teleman. Selective 40S footprinting reveals cap-tethered ribosome scanning in human cells. Mol. Cell, 79, 2020.

[68] Daria Shcherbakova and Vladislav V Verkhusha. Near-infrared fluorescent proteins for multicolor *in vivo* imaging. Nat Methods, 10:751–754, 2013.

[69] Mikhail E Matlashov, Daria Shcherbakova, Jonatan Alvelid, Mikhail Baloban, Francesca Pennacchietti, Anton A Shemetov, Ilaria Testa, and Vladislav V Verkhusha. A set of monomeric near-infrared fluorescent proteins for multicolor imaging across scales. Nature Comm, 239, 2020.

[70] Valentina Diehl, Martin Wegner, Paolo Grumati, Koraljka Husnjak, Simone Schaubeck, Andrea Gubas, Varun Jayeskumar Shah, Ibrahim H Polat, Felix Langschied, Cristian Prieto-Garcia, Kostantin Müller, Alkmini Kalousi, Ingo Ebersberger, Christian H Brandts, Ivan Dikic, and Manuel Kaulich. Minimized combinatorial CRISPR screens identify genetic interactions in autophagy. Nucleic Acids Res, 49, 2021.

[71] Nicholas T Ingolia, Sina Ghaemmagami, John R S Newman, and Jonathan S Weissman. Genome-wide analysis in vivo of translation with nucleotide resolution using ribosome profiling. Science, 324:218–223, 2009.

[72] Nicholas J McGlincy and Nicholas T Ingolia. Transcriptome-wide measurement of translation by ribosome profiling. Methods, 126:112–129, 2017.

[73] Lucas Philippe, Antonia MG van den Elzen, Maegan J Watson, and Carson C Thoreen. Global analysis of LARP1 translation targets reveals tunable and dynamic features of 5’ TOP motifs. Proc. Natl. Acad. Sci., 117:5319–5328, 2020.

[74] Amy E O’Connell, Maxim V Gerashchenko, Marie-Francoise O’Donohue, Samantha M Rosen, Eric Huntzinger, Diane Gleeson, Antonella Galli, Edward Ryder, Siqi Cao, Quinn Murphy, Shideh Kazerounian, Sarah U Morton, Klaus Schmitz-Abe, Vadim N Gladyshev, Pierre-Emmanuel Gleizes, Bertrand Séraphin, and Pankaj B Agrawal. Mammalian hbs1l deficiency causes congenital anomalies and developmental delay associated with pelota depletion and 80s monosome accumulation. PLoS Genetics, 2019.

[75] Markus Terrey, Scott I Adamson, Jeffrey H Chuang, and Susan L Ackerman. Defects in translation-dependent quality control pathways lead to convergent molecular and neurodevelopmental pathology. eLife, 10, 2021.

[76] Andrew C Hsieh, Yi Liu, Merritt P Edlind, Nicholas T Ingolia, Matthew R Janes, Annie Sher, Evan Y Shi, Craig R Stumpf, Carly Christensen, Michael J Bonham, et al. The translational landscape of mtor signalling steers cancer initiation and metastasis. Nature, 485(7396):55–61, 2012.

[77] Carson C Thoreen, Lynne Chantranupong, Heather R Keys, Tim Wang, Nathanael S Gray, and David M Sabatini. A unifying model for mTORC1-mediated regulation of mRNA translation. Nature, 485:109–113, 2012.

[78] Shintaro Iwasaki, Stephen N Floor, and Nicholas T Ingolia. Rocaglates convert dead-box protein eif4a into a sequence-selective translational repressor. Nature, 534(7608):558–561, 2016.

[79] Hema Manjunath, He Zhang, Frederick Rehfeld, Jaeil Han, Tsung-Cheng Chang, and Joshua T Mendell. Suppression of ribosomal pausing by eif5a is necessary to maintain the fidelity of start codon selection. Cell Rep, 29:3134–3146, 2019.

[80] Arya Vindu, Byung-Sik Shin, Kevin Choi, Eric T Christenson, Ivaylo P Ivanov, Chune Cao, Anirban Banerjee, and Thomas E Dever. Translational autoregulation of the *S. cerevisiae* high-affinity polyamine transporter hol1. Mol. Cell, 81:1–15, 2021.

[81] Eric W Mills and Rachel Green. Ribosomopathies: There’s strength in numbers. Science, 538, 2017.

[82] Xu Jiang, Shan Feng, Yuling Chen, Yun Feng, and Haiteng Deng. Proteomic analysis of mTOR inhibition-mediated phosphorylation changes in ribosomal proteins and eukaryotic translation initiation factors. Protein Cell, 7:533–537, 2016.

[83] Pu Zhang, Junko Iwasaki-Arai, Hiromi Iwasaki, Maris L Fenyus, Tajhal Dayaram, Bronwyn M Owens, Hirokazu Shigematsu, Elena Levantini, Claudia S Huettner, Julie A Lekstrom-Himes, et al. Enhancement of hematopoietic stem cell repopulating capacity and self-renewal in the absence of the transcription factor c/ebp*α*. Immunity, 21(6):853–863, 2004.

[84] B Niebuhr, G B Iwanski, S Roscher, C Stocking, and J Cammenga. Investigation of C/CEBP*α* function in humans (versus murine) myelopoiesis provides novel insight into the impact of *CEBPA* mutation in acute myelogenous leukemia (AML. Leukemia, 23:978–983, 2008.

[85] Derek L Stirewalt, Soheil Meshinchi, Kenneth J Kopecky, Wenhong Fan, Era L Pogosova-Agadjanyan, Julia H Engel, Michelle R Cronk, Kathleen Shannon Dorcy, Amy R McQuary, David Hockenbery, et al. Identification of genes with abnormal expression changes in acute myeloid leukemia. Genes, Chromosomes and Cancer, 47(1):8–20, 2008.

[86] Daniel G Gibson, Lei Young, Ray-Yuan Chuang, J Craig Venter, Clyde A Hutchison III, and Hamilton O Smith. Enzymatic assembly of DNA molecules up to several hundred kilobases. Nat Methods, 6:343–345, 2009.

[87] Balazs Halmos, Claudia S Huettner, Olivier Kocher, Katalin Ferenczi, Daniel D Karp, and Daniel G Tenen. Down-regulation and antiproliferative role of C/EBPalpha in lung cancer. Cancer Res., 62:528–34, 2002.

[88] Prashant Mali, Luhan Yang, Kevin M Esvelt, John Aach, Marc Guell, James E DiCarlo, Julie E Norville, and George M Church. RNA-guided human genome engineering via cas9. Science, 339:823–826, 2013.

[89] Eric Kowarz, Denise Löscher, and Rolf Marschalek. Optimized Sleeping Beauty transposons rapidly generate stable transgenic cell lines. Biotechnol. J., 10:647–653, 2015.

[90] Marcel Martin. Cutadapt removes adapter sequences from high-throughput sequencing reads. EMBnet.journal, 17:10–12, 2011.

[91] Wei Li, Tengfei Xiao, Le Cong, Michael I Love, Feng Zhang, Rafael A Irizarry, Jun S Liu, Myles Brown, and Shirley X Liu. MAGeCK enables robust identification of essential genes from genome-scale CRISPR/Cas9 knockout screens. Genome Biol, 15, 2009.

[92] Binbin Wang, Mei Wang, Wubing Zhang, Tengfei Xiao, Chen-Hao Chen, Alexander Wu, Feizhen Wu, Nicole Traugh, Xiaoqing Wang, Li Ziyi, Shenglin Mei, Yingbo Cui, Sailing Shi, Jesse Jonathan Lipp, Matthias Hinterndorfer, Johannes Zuber, Myles Brown, Wei Li, and Shirley X Liu. Integrative analysis of pooled crispr genetic screens using MAGeCKFlute. Nat Protoc, 14:756–780, 2019.

[93] Michael I Love, Wolfgang Huber, and Simon Anders. Moderated estimation of fold change and dispersion for RNA-seq data with DESeq2. Genome Biol, 15, 2014.

[94] Huaiyu Mi, Anushya Muruganujan, Dustin Ebert, Xiaosong Huang, and Paul D Thomas. PANTHER version 14: more genomes, a new PANTHER GO-slim and improvements in enrichment analysis tools. Nucleic Acids Res, 47:D419–D426, 2019.

[95] The Gene Ontology Consortium. The Gene Ontology resource: enriching a GOld mine. Nucleic Acids Res, 49:D325–D334, 2020.

[96] Michael Ashburner, Catherine A Ball, Judith A Blake, David Botstein, Heather Butler, J Michael Cherry, Allan P Davis, Kara Dolinski, Selina S Dwight, Janan T Eppig, Midori A Harris, David P Hill, Laurie Issel-Tarver, Andrew Kasarskis, Suzanna Lewis, John C Matese, Joel E Richardson, Martin Ringwald, Gerald M Rubin, and Gavin Sherlock. Gene Ontology: tool for the unification of biology. Nat Genet, 25:25–29, 2000.

[97] Ben Langmead and Steven L Salzberg. Fast gapped-read alignment with Bowtie2. Nat Methods, 9:357–359, 2012.

[98] Alexander Dobin, Carrie A Davis, Felix Schlesinger, Jorg Drenkow, Chris Zaleski, Sonali Jha, Philippe Batut, Mark Chaisson, and Thomas R Gingeras. STAR ultrafast universal RNA-seq aligner. Bioinformatics, 29:15–21, 2013.

[99] Bo Li and Colin N Dewey. RSEM accurate transcript quantification from RNA-Seq data with or without a reference genome. Bioinformatics, 12, 2011.

